# Structural bases for Nuclear Factor 1-X activation and DNA recognition. Prototypic insight into the NFI transcription factor family

**DOI:** 10.1101/2025.03.21.644642

**Authors:** Michele Tiberi, Michela Lapi, Louise Jane Gourlay, Antonio Chaves-Sanjuan, Maurizio Polentarutti, Nicola Demitri, Miriam Cavinato, Diane Marie Valérie Jeanne Bonnet, Valentina Taglietti, Anna Righetti, Rachele Sala, Silvia Cauteruccio, Amit Kumawat, Rosaria Russo, Alberto Giuseppe Barbiroli, Nerina Gnesutta, Carlo Camilloni, Martino Bolognesi, Graziella Messina, Marco Nardini

## Abstract

Nuclear Factor I (NFI) proteins were first identified in adenovirus DNA replication and later as regulators of gene transcription, stem cell proliferation, and differentiation. They play key roles in development, cancer and congenital disorders. Within the NFI family, NFI-X is critical for neural stem cell biology, hematopoiesis, muscle development, muscular dystrophies and oncogenesis. Here, we present the first structural characterization of the NFI transcription factor, NFI-X, both alone and bound to its consensus palindromic DNA site. Our analyses reveal a novel, MH1-like fold within NFI-X DNA-binding domain (DBD) and identify crucial structural determinants for activity, such as a Zn²⁺ binding site, dimeric assembly, activation mechanism and DNA-binding specificity. Given the >95% sequence identity within the NFI DBDs, our structural data are prototypic for the entire family; a NFI Rosetta Stone that allows decoding a wealth of biochemical and functional data and provides a precise target for drug design in a wider disease context.

## Introduction

Nuclear factor 1 X (NFI-X) is a member of the NFI family, ubiquitous group of transcription factors (TF) that includes three additional NFI-encoding genes (*nfia*, *nfib*, and *nfic*). NFIs are site-specific DNA-binding proteins that govern many aspects of cell biology, regulating both viral DNA replication and cellular gene expression [1]. Human and vertebrate NFI-X genes contain 11 exons and alternative splicing gives rise to different, tissue-specific and developmental stage isoforms [2]. In gene regulation, NFIs determine stem cell proliferation and differentiation, being highly expressed in embryonic and adult stem cells in many tissues [3–5]. Consequently, NFIs are implicated in several developmental disorders [6–9], and act as tumor suppressors in some cancers [10].

To date, NFI research has focused particularly on NFI-X due to its essential role in multiple organ systems, including neural stem cell biology, hematopoiesis and muscle development [4]. NFI-X is expressed throughout the brain and is implicated in nervous system development throughout embryogenesis and in neuronal stem cell homeostasis [4,11]. In muscle, NFI-X plays a pivotal role in skeletal muscle development, by regulating fetal-specific myogenic program and fiber type properties [3], and in post-natal regeneration, by regulating muscle stem cells differentiation upon acute damage [12,13]. In this context, NFI-X has been studied in the pathophysiology of muscular dystrophies [14], based on evidence that its absence in *nfix* null mice protects from disease progression [13,15,16].

Furthermore, NFI-X is implicated in several human cancers, and two rare congenital malformation syndromes: Malan syndrome (MALNS) and Marshall-Smith Syndrome (MSS), caused by mutations that cluster in distinct regions of the *nfix* gene [4,6,7]. Respectively, MALNS and MSS mutations cluster in the DNA-binding domain (DBD) and the regulatory C-terminal transactivation domain (TAD)[7].

The molecular mechanisms of NFIs are unknown and debated, mostly due to the difficulties in producing sufficient heterologous protein yields for *in vitro* characterization. NFIs bind as dimers to the palindromic consensus sequence TTGGC(n5)GCCAA, in double stranded DNA, with nanomolar affinities [17–19]. Indeed, dimerization is an integral part of NFI DNA-binding functions, with protein mutants devoid in dimerization ability, unable to properly bind DNA [20]. Interestingly, NFI monomers can also bind to individual half-sites, but with two orders of magnitude lower affinity [18]; shortening the spacer region by one base-pair lowers the affinity to a value similar to that observed with a single half-site, implying that a shorter spacer impacts on the simultaneous interaction of NFI monomers with both half-site sequences [21]. Additionally, mutations out with the consensus site induce minor differences in NFI binding affinity [22].

Studies on truncated NFI-C mutants, show that the first 240 N-terminal amino acids appears to be sufficient for dimerization and DNA binding [23,24], with the dimerization domain being probably housed within amino acids 170-240 [25,26]. The NFI DBDs are highly similar (85% sequence identity and 92% similarity) (**Fig. S1**), and conserved residues include four cysteines (named Cys-2, Cys-3, Cys-4, and Cys-5) and all, except Cys-3, are essential for DNA-binding. Cys-3 oxidation can lead to reversible inactivation of the protein [22,27,28], thus, cysteine residues play important roles in different aspects of NFI-DNA interactions, including redox regulation [23,28].

In addition to the DBD, NFIs contain a highly divergent C-terminal TAD region (**Fig. S1**), which is the target of other transcriptional coregulators [24,29] and is predicted to be largely intrinsically-disordered by Alphafold [30]. Despite poor sequence conservation, some features are shared, such as high proline content and the presence of post-translational modification sites that also contribute to NFI regulation [31,32]. The C-terminus of NFI-X from *Xenopus laevis* has been reported to negatively regulate NFI-X function by preventing DNA binding *in vitro* [29].

Despite general notions on NFI function, precise mechanistic details are lacking. Given the significant role of NFI-X in development and disease, understanding its structure is crucial to develop targeted approaches to inhibit or promote its function. To this aim, we structurally characterized two recombinant protein constructs (NFI-X^176^ and NFI-X^193^) encompassing the DBD of NFI-X isoform 2, which represents the most highly expressed splicing variant in mouse skeletal muscle, with full sequence identity with its human ortholog. Our analyses unravel the novel DBD fold, and reveal for the first time, the structural determinants of dimerization, activation, and DNA-binding specificity, and provide insight into the impact of NFI-X pathological mutations. Given the strong sequence conservation within the NFI family, we propose our NFI-X DBD structure as prototypes for the entire NFI family.

## Results

### The NFI-X DBD is a monomer in solution

Bioinformatics analyses of full-length NFI-X isoform 2 using Pfam, GlobPlot and DisEMBL [33–35] guided the design of two protein constructs encompassing the N-terminal DBD for heterologous expression in bacterial cells (**Fig. S1**). NFI-X^176^ (residues 14–176) represents the minimal predicted DBD, while NFI-X^193^ includes 17 additional C-terminal residues. The first N-terminal 13 amino acids were omitted in both constructs due to predicted structural disorder and their lack of relevance for DNA binding [36]. Both NFI-X^176^ and NFI-X^193^ were successfully expressed in soluble form in BL21 Shuffle T7 *E. coli* cells, albeit with different yields: 10 mg/L and 2 mg/L bacterial culture, respectively.

Both protein constructs are monomers in solution, as shown by size exclusion chromatography (SEC) and migrate by SDS-PAGE with molecular weights (MWs) in line with their calculated values of 19.5 kDa (NFI-X^176^) and 21.4 kDa (NFI-X^193^), respectively (**Fig. S2A, S2B**). Thermal denaturation profiles, following the loss of helical structures with increasing temperature using circular dichroism (CD) at 222 nm, indicate that both constructs are stable, and show similar melting temperature (T_M_) values of 55.6 °C and 56.2 °C for NFI-X^176^ and NFI-X^193^, respectively (**Fig. S2C**).

### The C-terminus of NFI-X DBD increases DNA-binding affinity

NFI-X^176^ and NFI-X^193^ constructs were tested for DNA binding by electromobility shift assay (EMSA), using an optimized, 31bp DNA probe, containing the consensus NFI palindromic dyad sequence TTGGC(n5)GCCAA (named *NFI-31bp*; see Methods). Comparative EMSAs indicate that both constructs bind DNA in a dose-dependent manner, but NFI-X^193^ binds with ∼10-fold higher affinity than NFI-X^176^ (**Fig. 1A**). The presence of faster migrating complexes in the NFI-X^176^ binding reactions are further indicative of reduced stability of the DNA assembly. This result indicates that the 17 residues C-terminal extension is critical for high affinity DNA-binding of the NFI-X DBD. Therefore, all further EMSAs were carried out on NFI-X^193^.

**Fig. 1.**
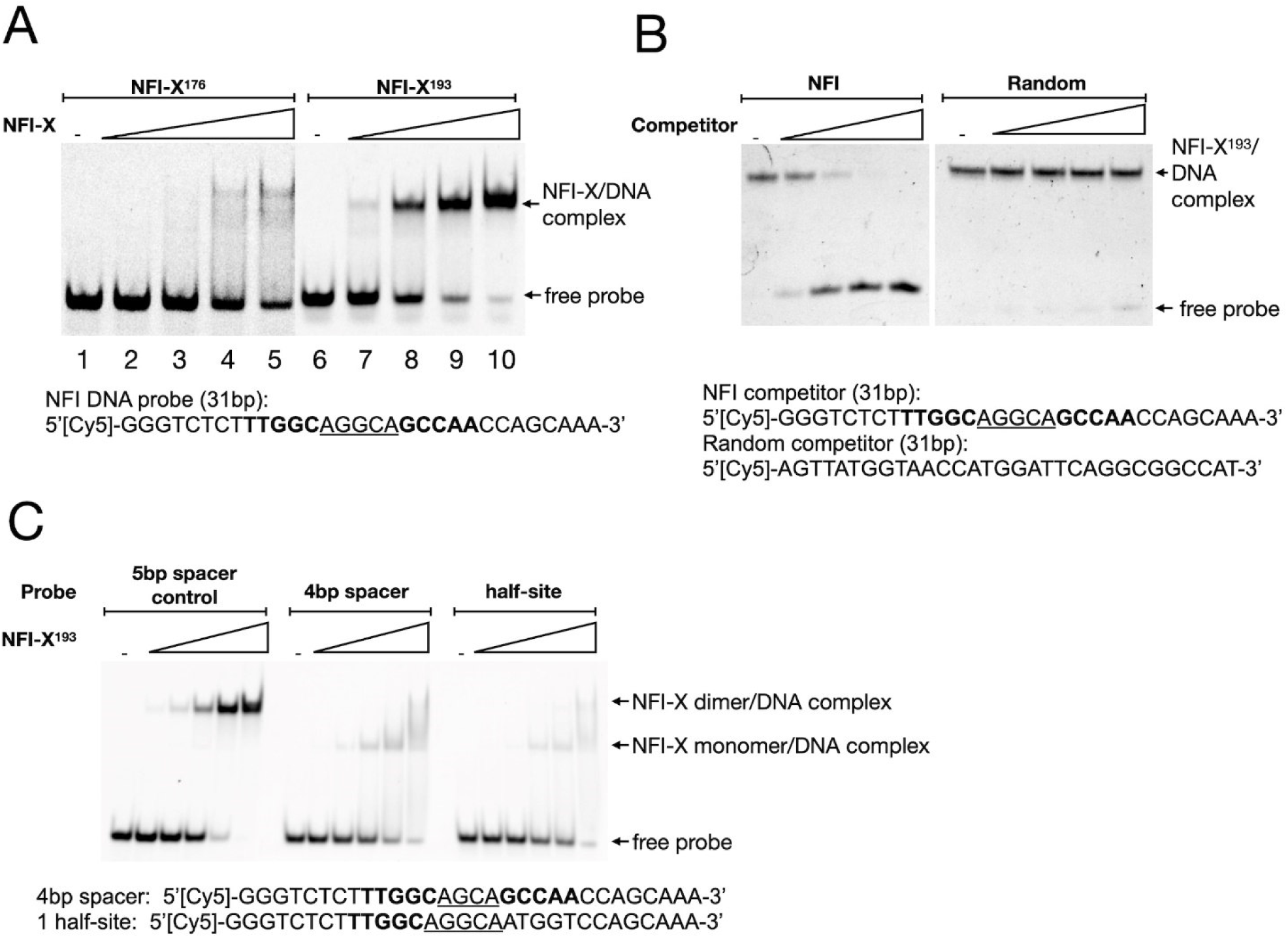
NFI-X DBD binding to DNA. **A)** Dose-response EMSA experiments were carried out with NFI-X^176^ (Lanes 2 to 5) and NFI-X^193^ (Lanes 7-10), at increasing protein concentrations (30 nM, 90 nM, 270 nM, 0.8 µM) and 20 nM of Cy5-labeled 31 bp dsDNA probe (sequence shown), presenting the NFI consensus sequence (bold font) and a 5 bp spacer (underlined). Bands corresponding to the free probe or the NFI-X/DNA complex are indicated. Lanes 1 and 6 represent the control reactions in the absence of protein addition. **B)** Competition EMSA experiments to demonstrate NFI binding site specificity by NFI-X DBD. NFI-X^193^ (20 nM) was incubated with the probe (20 nM) alone (-) or in the presence of increasing concentrations (6.25X, 12.5X, 25X and 50X respect to the DNA probe) of unlabelled 31 bp NFI competitor (left panel) or unlabelled 31 bp random sequence (right panel) as described in the Methods section; **C)** EMSA experiments to demonstrate the requirement for a 5 bp spacer and to illustrate binding as dimer. NFI-X^193^ was incubated in the presence of the Cy5-labeled 31 bp dsDNA probe containing a palindromic 5 bp spacer consensus site (left panel) as a positive control, a 4 bp spacer (central panel) palindromic site, or a 31 bp dsDNA probe containing one half site (right panel). In all panels, protein concentrations were 0.5X, 1X, 2X, 4X and 8X the concentration of the probe (20 nM). Reactions in the absence of protein addition are indicated (-). The 4 bp spacer and half-site probe sequences are shown below, with the NFI consensus sequence (bold font) and spacer regions (underlined).

DNA-binding is sequence-specific, as shown by competition EMSA experiments (**Fig. 1B**), with two NFI-X monomers bound to DNA, as deduced from comparisons made with a DNA probe, containing a single half-site that shows a shifted complex with higher mobility, representing a single DNA-bound NFI-X^193^ monomer. This band shift is similar to that observed when the spacing between the two half sites is shortened by one base-pair (n4) (**Fig. 1C**), indicating that spacers <5bp sterically impair dimer formation on DNA (see below), leading to lower stability of the complex and single-site monomer binding.

### NFI-X DBD has a novel MH1-like fold and contains a conserved Zn^2+^-binding site

Considering its higher production yield, the NFI-X^176^ construct was selected for crystallization trials, in complex with DNA. Crystals suitable for X-ray diffraction experiments were only obtained with a 23bp DNA (*NFI-23bp*; see Methods) (**Fig. S3**). A sequence-based search for NFI-X DBD structural homologs identified only the MH1 domain of Smad proteins, however with low (< 20%) sequence identity. Nevertheless, a clear sequence match was identified between the Smad CCCH motif, responsible for Zn^2+^ coordination, and NFI-X Cys103, Cys156, Cys162, and His167 residues (**Fig. 2A**). Strikingly, a Zn^2+^ ion was identified to be bound to NFI-X^176^ by in-solution Flame Atomic Absorption Spectrometry (FAAS) (**Fig. S4A**). Taking advantage of the bound Zn^2+^, the crystal structure of NFI-X^176^ (PDB: 7QQD) was solved at 2.7 Å by SAD-phasing (**Fig. S4B and Table S1**). Since zinc ions were not added during sample preparation, they are likely to have been picked from the growth media during protein expression.

**Fig. 2.**
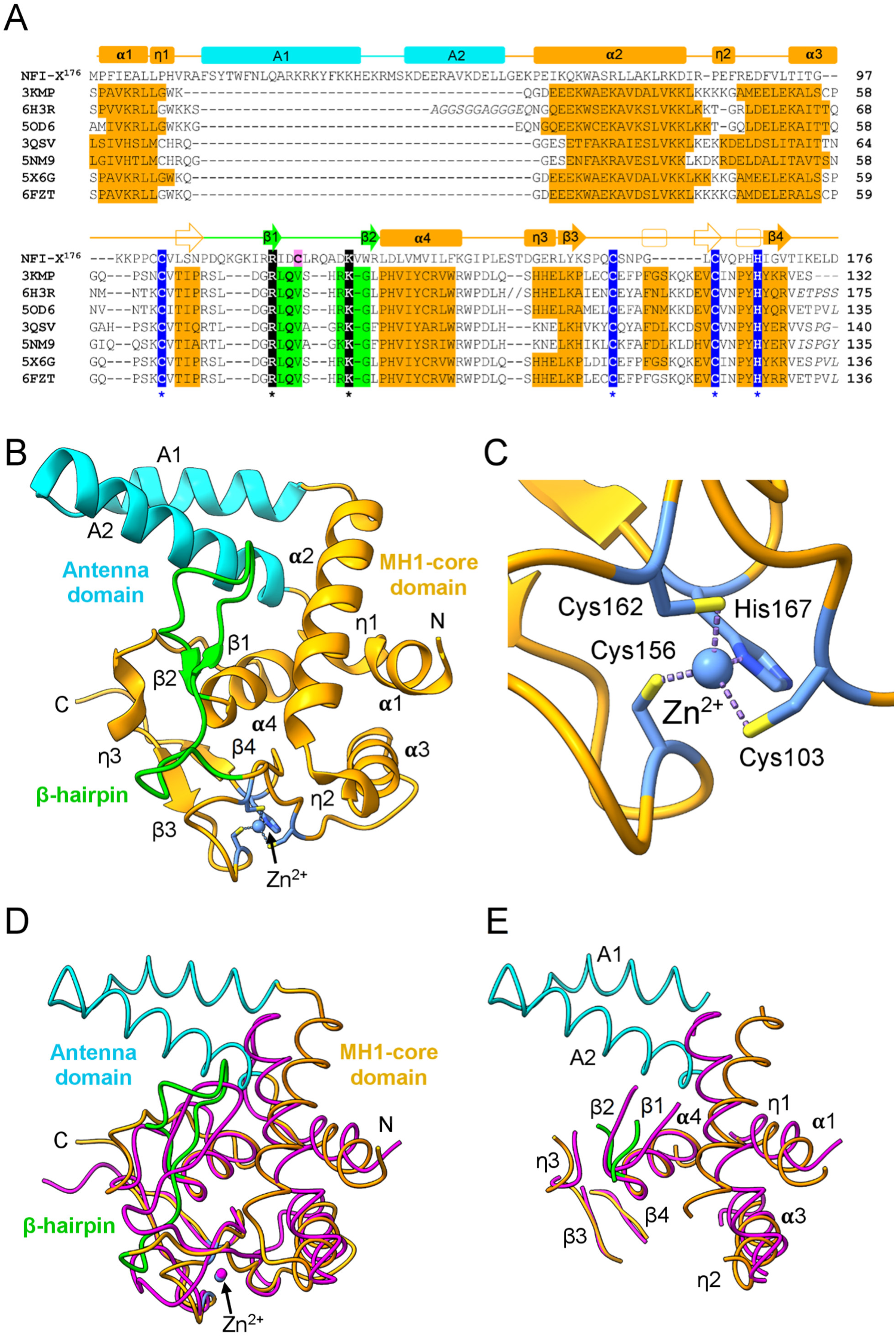
The crystal structure of NFI-X^176^ and sequence/structure homology with the Smad protein MH1 domain. **A)** Structure-based sequence alignment of NFI-X^176^ with related Smad MH1 domains using the MUSCLE program (https://www.ebi.ac.uk/Tools/msa/muscle/) and manually corrected, based on 3D structure comparisons [44]. NFI-X^176^ was aligned with human Smad2 (PDB: 6H3R [45]), Smad3 (PDB: 5OD6 [46]) and Smad9 (PDB: 6FZT [45]), mouse Smad1 (PDB: 3KMP [47], Smad4 (PDB: 3QSV (unpublished), and Smad5 (PDB: 5X6G [48]), and *Trichoplax adhaerens* Smad4 (PDB: 5NM9 [46]). NFI-X^176^ secondary structure elements are shown above the aligned proteins with the following color code: MH1-core domain in orange, antenna domain in cyan, β-hairpin in green. Smad residues that adopt α-helices and β-strands are shaded; Smad secondary structure elements not conserved in NFI-X are shown, transparent above the sequence. All secondary structure assignments were made using PDBsum [49]. Zn^2+^-coordination residues are shown in white bold letters, shaded in blue, and highlighted with an asterisk, while Smad residues that provide sequence-specificity, including conserved NFI-X residues Arg116 and Lys125, are shaded in black. A 30-residue insertion in Smad2 (6H3R) is indicated by //; Cys119 (Cys-3) is shaded in pink. **B)** Ribbon representation of the NFI-X^176^ crystal structure (PDB: 7QQD). The MH1 core domain and the antenna domain are shown in orange and cyan, respectively; both domains are labeled, as are the secondary structure elements and N-and C-termini. The β-hairpin is highlighted in green and the bound Zn^2+^ ion is shown as a light blue sphere, with the coordinating cysteine and histidine residues are shown as light blue sticks. **C)** Detailed view of Zn^2+^ ion binding site and residues of the CCCH motif, shown in sticks, colored by heteroatom and labeled; **D)** Structural superposition of NFI-X^176^ (colored as in panel B) with a representative Smad protein, human Smad3 (PDB: 5OD6; RMSD of 0.94 Å over 109 paired residues) shown in magenta; **E)** Structural superposition, as in panel D, with loops removed for clarity. Panels B to E were generated using ChimeraX version 1.8 [50].

There were two NFI-X^176^ molecules in the asymmetric unit (ASU) (**Table S1**); electron density was well-defined for amino acids 13 to 174 in both chains. The NFI-X DBD fold can be subdivided into two domains, indicated as “antenna” and “core”, which hosts a Zn^2+^-binding site (**Fig. 2B and Fig. 2C**). The core domain is formed by a four α-helical bundle (α1, α2, α3 and α4), three 3^10^ helices (η1-3) and two short, anti-parallel β-sheets (β1–β2, β3–β4). Protein structure comparisons, made using the DALI server [37], pinpoint the MH1 domain of Smad TFs, as the only structural homolog of the NFI-X core domain. Despite poor sequence identity, the MH1 α-helical bundle of the NFI-X DBD superimposes well with that of Smad TFs, with some differences in the relative orientation of the helices (RMSD ∼2.1 Å), and the structural match further improves (RMSD ∼1.8 Å) if the loops connecting secondary structure elements are removed (**Fig. 2D and Fig. 2E**). The smaller “antenna” domain (not present in Smad TFs) is a helical excursion, formed by A1 and A2, protruding from the core domain, nestled between α1 and α2 helices (**Fig. 2A and Fig. 2B**). Therefore, overall, the NFI-X DBD presents a novel “MH1-like” fold that, given its extensive sequence conservation in NFIs (**Fig. S1**), can be considered as a DBD prototype for the entire NFI family.

With regards to the conserved CCCH Zn^2+^ coordination site, in NFI-X^176^ Zn^2+^ is coordinated by Cys103, Cys156, Cys162 and His167. Cys103 pertains to a flexible loop (residues 97-115) connecting α3 and β1, Cys156 and Cys162 belong to a different loop (residues 153-164) that connects strands β3 and β4, and His167 is contributed from 3^10^ helix η3 (**Fig. 2A, and Fig. 2C**). Given the invariance of these coordination residues in the NFI family, corresponding to Cys-2, Cys-4, and Cys-5 (see Introduction) (**Fig. S1**), our data suggests a conservation of the Zn^2+^-binding site within the NFI family in general. Indeed, mutation of these cysteine residues in NFIs has been shown to prevent DNA binding [36]. Our structure rationalizes such data by identifying the Zn^2+^ binding site as a key factor for the correct folding and stabilization of the NFI-X^176^ core domain, with implications on the C-terminal region of the DBD, crucial for dimerization and efficient DNA-binding, as discussed below.

Other than the antenna domain, a second clear difference between the NFI-X^176^ core domain and the Smad MH1 domain concerns the β-hairpin (strands β1 and β2), exploited by Smad TFs to bind DNA in a sequence specific manner at the major groove (**Fig. S5**). Despite a similar 3D location, the NFI-X^176^ β-hairpin (strands β1 and β2) is three-residues longer (and also the preceding loop is three-residues longer), the two connected β strands are shorter (only two residues each, instead of three), and there is no amino acid conservation relative to Smad TFs, except for NFI-X^176^ Arg116 and Lys125 residues that provide sequence-specific contacts in Smad TFs (**Fig. 2A, and Fig. S5**). Differences at the β-hairpin region are not unexpected, considering the different DNA-binding modes of the two TF families. Indeed, Smad TFs recognize a 4bp palindromic GTCT-AGAC motif [38] with no spacing between half-sites, thus the two MH1 domains position themselves at opposite sides of the DNA duplex with no physical interactions between monomers (**Fig. S5**). On the contrary, NFI-X DBD binds at a palindromic 5bp dyad sequence TTGGC(n5)GCCAA, with the spacer region (n5) expected to position the two DBD core domains on the same side of the DNA (see below).

In line with SEC studies, indicating NFI-X^176^ is monomeric in solution (**Fig. S2A**), NFI-X^176^ is a monomer in the crystal and the two NFI-X^176^ chains in the ASU (**Fig. S6A**) do not form a biologically relevant dimer, as deduced using jsPISA [39]. Despite co-crystallization with DNA, in the crystal (PDB: 7QQD) no DNA was found bound to NFI-X^176^. Residual electron density, compatible with DNA, however, was present at the boundaries between different unit cells, although only 15 disordered base pairs could be traced in protein packing voids, with loose DNA-protein contacts (**Fig. S6B**). This result agrees with EMSA experiments that show that NFI-X^176^ has reduced affinity and stability for DNA (**Fig. 1A**) and that *NFI-23bp* is too short to be bound with high affinity (**Fig. S3**). Nevertheless, the presence of the DNA as a “crystallization additive” promoted crystallization and stabilized the crystal by filling, otherwise empty, volumes among unit cells **(Fig. S6B)**. The use of DNA as a crystallization additive, in fact, has been previously reported for lysozyme [40].

### NFI-X DBD binds as a dimer to the DNA major groove, stabilized through a C-terminal swapping mechanism

To solve the structure of NFI-X DBD in complex with DNA, efforts focused on the NFI-X^193^ construct, based on its improved affinity for DNA with respect to NFI-X^176^ (**Fig. 1A**), using *NFI-31bp* demonstrated to be the optimal length for efficient binding (**Fig. S3**). Since the yield of NFI-X^193^ was insufficient (0.6 mg/L bacterial culture) for protein crystallization, a cryo-EM approach was adopted, despite the small size of the protein/DNA complex (approx. 70 kDa). Nevertheless, the cryo-EM structure of NFI-X^193^/DNA could be solved at 3.86 Å resolution (PDB: 9QKY, EMD-53223; **Table S2)**. In fact, under grid vitrification conditions (see Methods), the purified complex produced long straight fibrils (**Fig. 3A and Fig. S7**) composed of a right-handed superhelix made of a repeating unit formed by a NFI-X^193^ dimer bound to *NFI-31bp*. Within the fibril, the DNA molecules interact in an end-to-end manner with an angle of about 140° between the axis of the DNA molecules (**Fig. 3A and Fig. 3B**). Cryo-EM electron density was interpretable up to residue Tyr181 for both protomers (A and B) of the dimer, suggesting that the final 12 residues of NFI-X^193^ are structurally disordered.

**Fig. 3.**
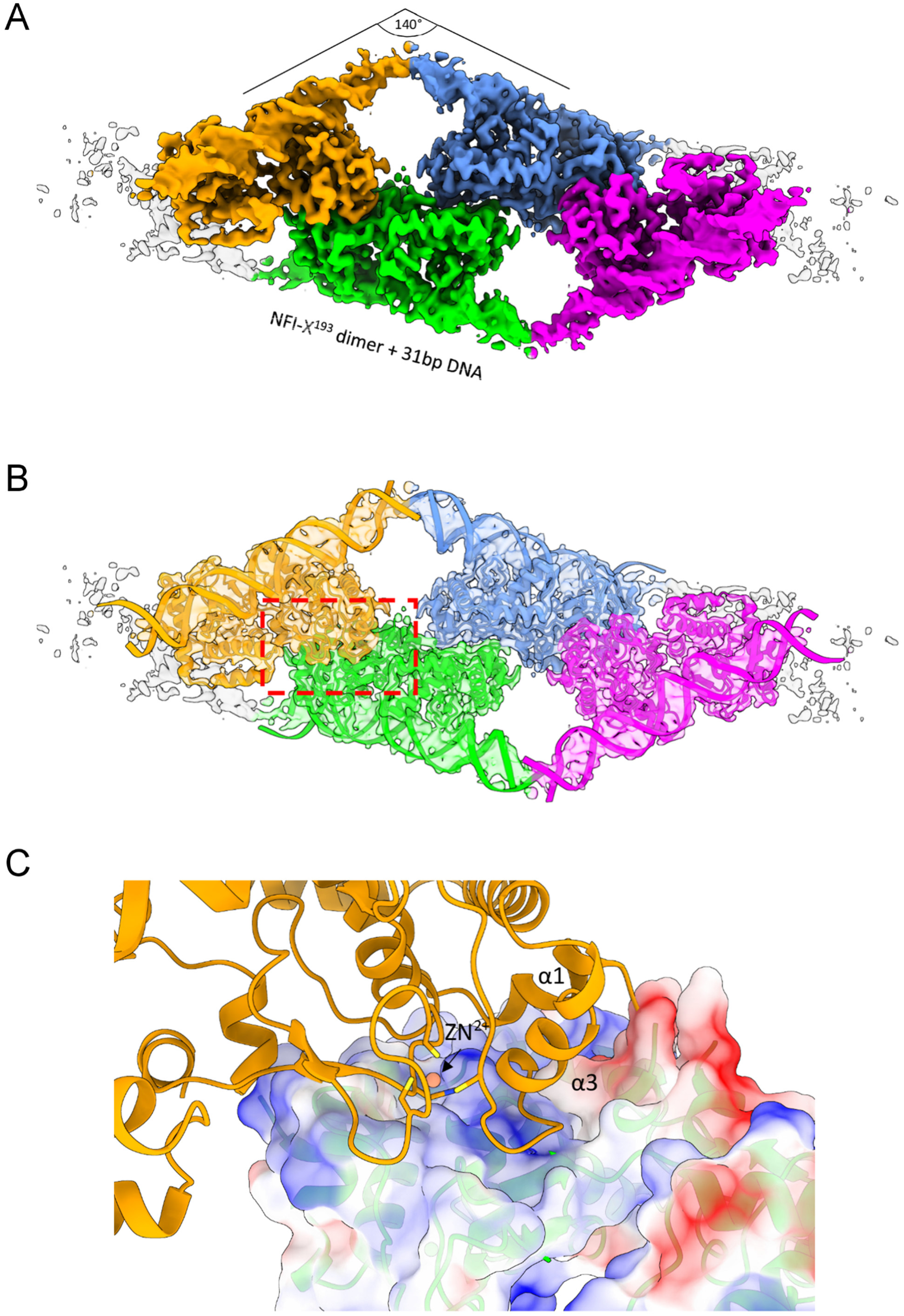
Cryo-EM structure of the NFI-X^193^/DNA fiber. **A)** Cryo-EM map for four consecutive NFI-X^193^ dimer units of the fibril, shown in orange, green, blue, and magenta. The angle between the axes of two adjacent DNA molecules is indicated; **B)** Model of the NFI-X^193^/DNA complex fitted into the cryo-EM map (transparent); **C**) Detailed view of the protein-protein interface between adjacent NFI-X^193^ dimers (yellow and green) within the fiber, indicated by the square with red dashes in panel B. To highlight interface shape complementarity, one monomer (yellow dimer) is shown in cartoon representation, while the second monomer (green dimer) is shown with a transparent electrostatic surface. Relevant secondary structure elements and the Zn^2+^ binding site are labelled. This Figure was generated using ChimeraX version 1.8 [50].

The cryo-EM structure provides the first direct visualization of a NFI dimer bound to DNA (**Fig. 4A)**. Within the dimer, protein-protein contacts are limited and mediated by a swapping mechanism which brings the C-terminal helix of one subunit (residues 174-180), to be sandwiched between the same helix and loop 140-143 of the other subunit. Interactions are both hydrophobic, with Leu(A)175, Tyr(A)178 and Leu(A)179 of one subunit nestled in a pocket lined by Ile(B)140, Pro(B)141, Leu(B)142, Val(B)170 and Ile(B)172 of the other subunit, and electrostatic, between Lys(A)173 NZ and Glu(B)143 OE1 (**Fig. 4B**). The molecular architecture of the DNA-bound NFI-X^193^ dimer rationalizes why DNA binding affinity was reduced in the NFI-X^176^ construct, due to the absence of residues 177-193 which contain the C-terminal helix, involved in the dimerization swapping mechanism (**Fig. 1A**).

**Fig. 4.**
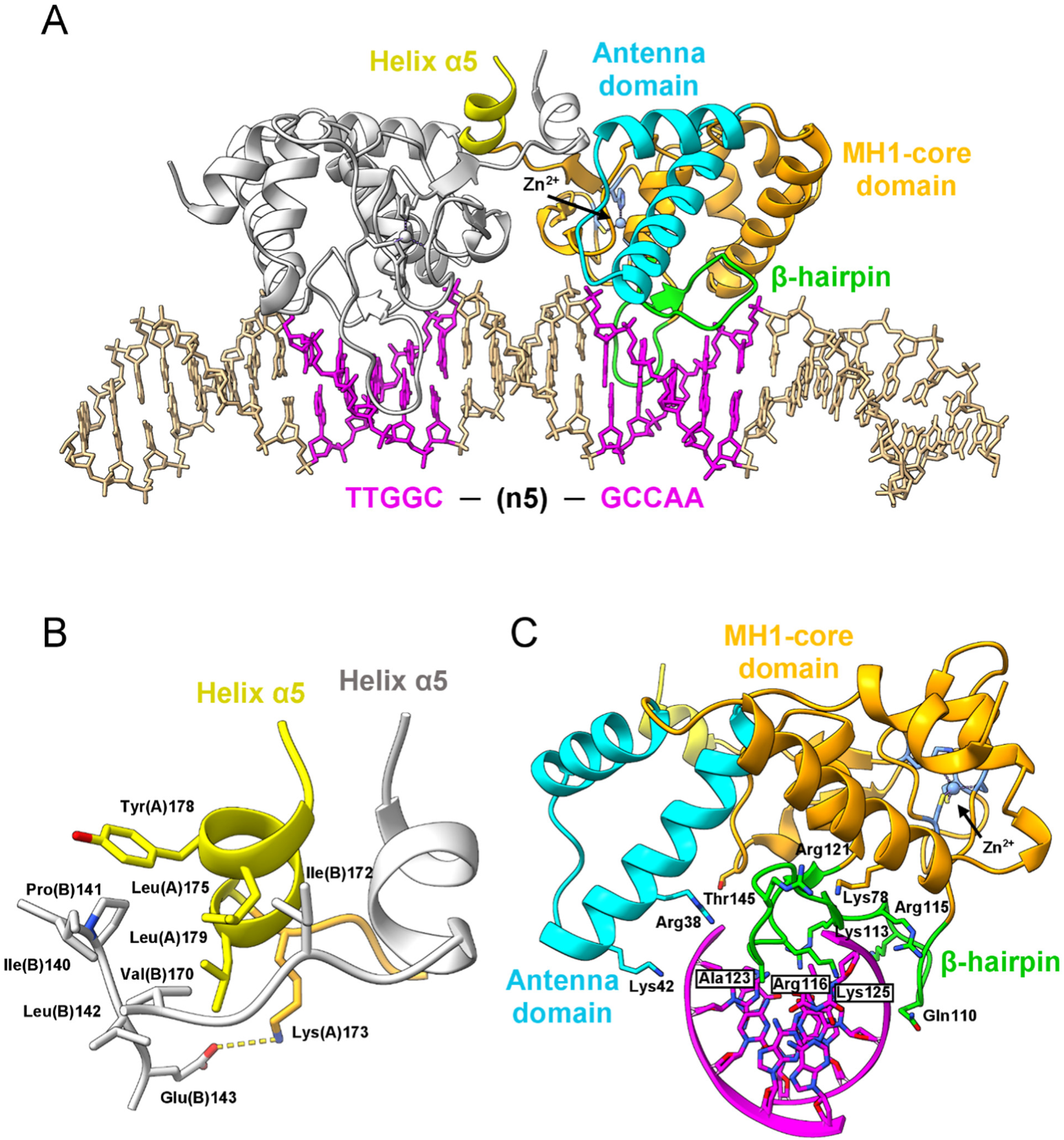
Cryo-EM structure of NFI-X^193^ in complex with DNA. **A)** NFI-X^193^ dimer bound to DNA. One monomer is shown with coloring as in Figure 2, while the other is shown in grey to highlight the dimer swapping mechanism involving helix α5 (yellow). The DNA molecule is shown in sticks (light brown color), with the consensus NFI-binding site in magenta. **B)** Detailed view of the dimerization interface involving the swapping of helix α5. Interface residues are shown as sticks and labelled. The salt-bridge between Lys173 (chain A) and Glu145 (chain B) is shown in dashes. **C)** Frontal view of one NFI-X^193^ monomer bound to DNA. Residues interacting with DNA are shown as sticks and labelled. For clarity, only one NFI half site is shown. This Figure was generated using ChimeraX version 1.8 [50].

### The antenna domain and the β-hairpin loop are the structural determinants of NFI-X DNA binding

NFI-X^193^ binds at the NFI consensus sequence, with the 5bp spacer positioning the two DBD core domains on the same side of the DNA (**Fig. 4A)**. Each NFI-X^193^ protomer interacts at each half site in a similar manner, with the antenna and core domains flanking the DNA helix on opposite sides (**Fig. 4C**). As expected, the NFI-X^193^ DNA interaction interface is rich in positively charged amino acids (**Fig. S8**). Detailed protein-DNA interactions are shown in **Figure 5**. In subunit A, Lys78, Gln110 and Arg115 bind to the DNA phosphate backbone atoms of T^9^ and G^11^ of the consensus sequence T^9^T^10^G^11^G^12^C^13^, respectively, while Arg121 interacts to with T^8^, immediately prior to the consensus motif. On the opposite strand (G^-13^C^-^ ^12^C^-11^A^-10^A^-9^), Arg38 and Lys 42 bind to the first two nucleotides (G^-13^, and C^-12^ respectively). Finally, Thr145 and Lys113 interact with T^-14^ and C^-15^ of the (n5) linker, respectively. Sequence-specific interactions are provided only by Arg116 and Lys125: Arg116 side-chain binds to G^12^ of the consensus motif and G^-13^ of the complementary strand; Lys125 side-chain interacts with G^11^G^12^, while its carbonyl oxygen atom is H-bonded to C^-12^ (**Fig. 5B**). Furthermore, the Ala123 side-chain contributes to sequence-specific recognition by facing the methyl groups of T^9^T^10^, providing favorable hydrophobic interactions. In subunit B, protein-DNA interactions are specularly conserved (**Fig. 5A**). Of the interacting residues, Arg38 and Lys42 are contributed by helix A2 of the antenna domain, whereas the remaining residues, apart from Lys78 and Thr145, belong to the β-hairpin loop (**Fig. S1**), whose convex face dives into the concave major groove of the duplex DNA containing 5bp of the consensus-binding motif (**Fig. 4A and Fig. 4C**).

**Fig. 5.**
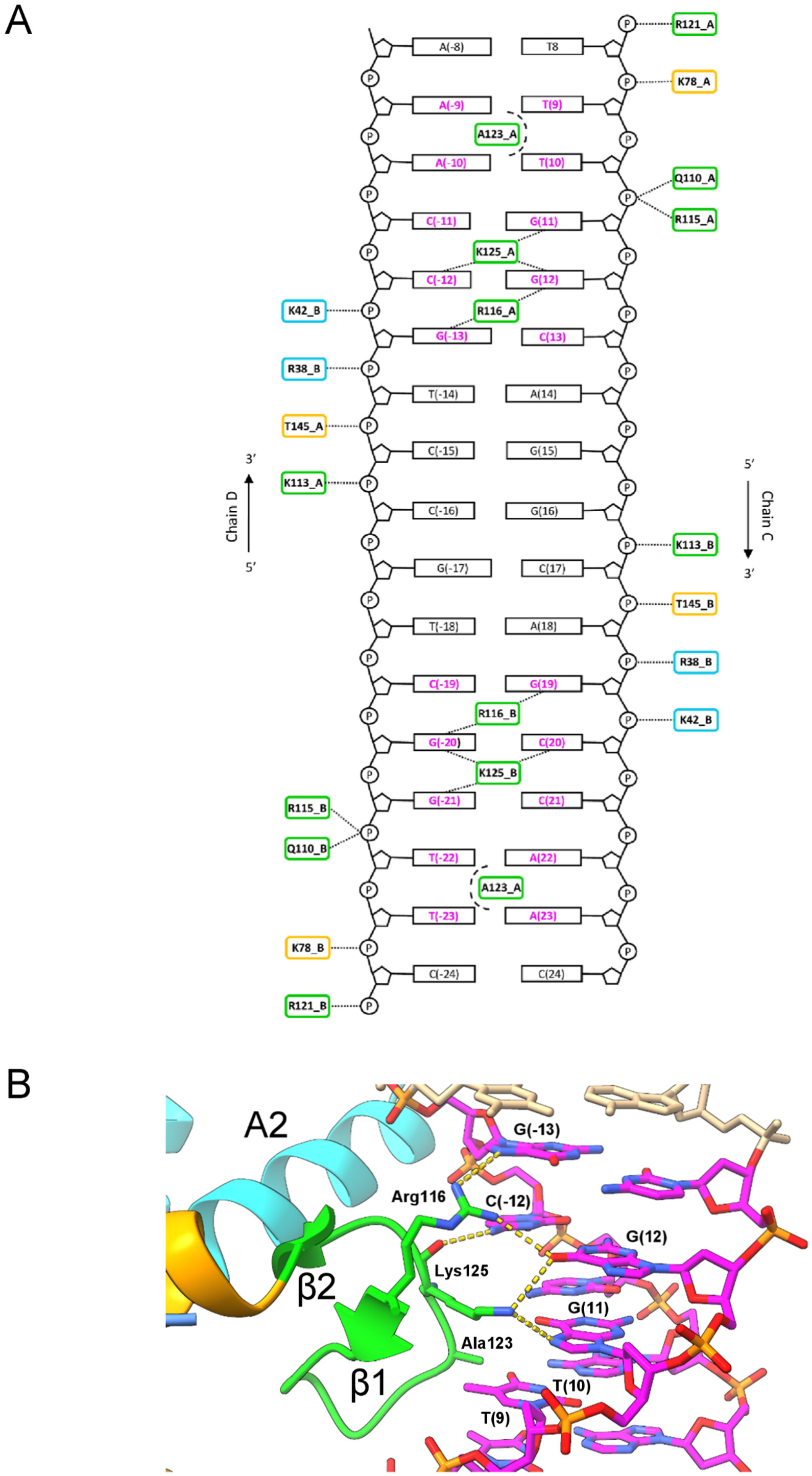
DNA contacts made by NFI-X DBD. **A)** Schematic view of the interactions between NFI-X^193^ residues and DNA: dotted lines indicate hydrogen bonds and salt bridges, dashed circular lines indicate hydrophobic sequence-specific interactions. NFI-X^193^ residues are boxed and colored as in Figure 4A. **B)** Site-specific interactions made by NFI-X^193^ Arg116 and Lys125. Hydrogen bonds are shown in dotted lines. Panels B was generated using ChimeraX version 1.8 [50].

Superposition of cryo-EM DNA-bound NFI-X^193^ with the NFI-X^176^ crystal structure shows a RMSD 0.95 Å over 159 aligned residues, indicating that NFI-X DBD binds its target DNA without large conformational changes, in particular regarding the relative position of the antenna and core domains (**Fig. 6A**). Only a minute difference is present which, however, explains the activation mechanism that triggers the insertion of the β-hairpin loop into the DNA major groove for base-reading. In the unbound state, the β-hairpin loop and the antenna domain are locked together by an intramolecular salt-bridge involving Asp124 and Arg38 side-chains, respectively. In the DNA-bound state, Arg38 side-chain changes its orientation, making a salt-bridge with a DNA phosphate group (G^-13^), thus unlocking the β-hairpin loop (containing Asp124) from the antenna domain (**Fig. 6B**). This structural change in turn allows Lys125 side-chain to move into the major groove to provide sequence specific DNA-binding interactions, together with Arg116 (**Fig. 5B)**.

**Fig. 6.**
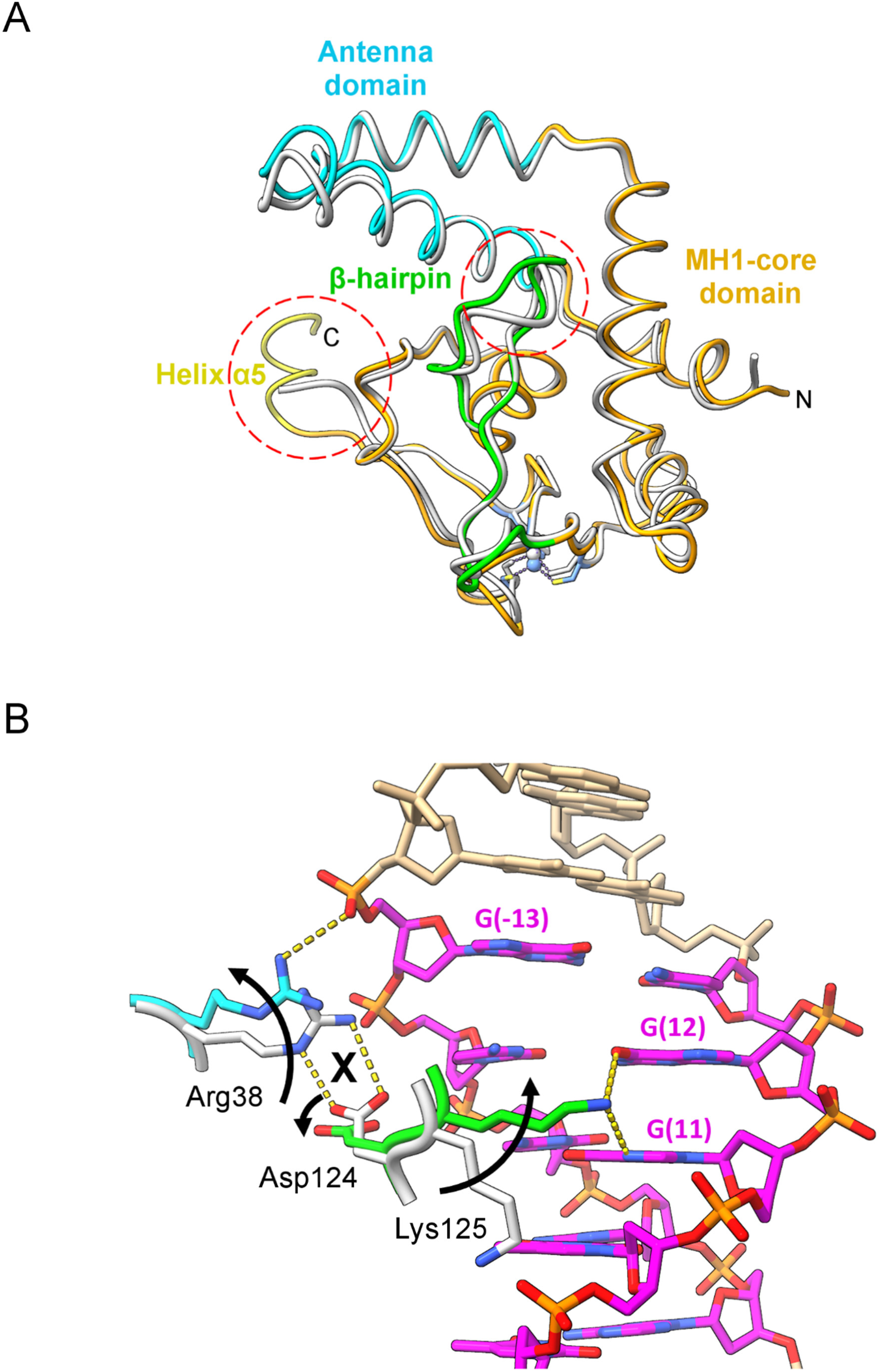
Activation mechanism of NFI-X DBD. **A)** Superimposition of the NFI-X^193^/DNA cryo-EM structure (colored as in Figure 2) and the crystal structure of NFI-X^176^ (grey). For clarity, DNA is not shown in the NFI-X^193^/DNA complex. Major structural differences are highlighted by red dashed circles. **B)** Structural changes associated with DNA-binding, highlighted by arrows. Residues relevant in the NFI-X DBD activation mechanism are shown as sticks and labelled. Salt-bridge and hydrogen bond interactions are shown as dashed lines. The Figure was generated using ChimeraX version 1.8 [50].

Interestingly, mutations at residues belonging to the antenna domain (Arg38) and to the β-hairpin loop (Lys113, Arg115, Arg116, Arg121, Lys125, and Arg128; see **Fig. S1**) cause a rare genetic disease called Malan syndrome that is characterized by excessive body growth, skeletal anomalies, and intellectual retardation [6,7]. In the DNA-bound NFI-X^193^ structure, all cited residues are either interacting with DNA bases (Lys125, and Arg116) or in contact (Lys113, Arg115, Arg121) or proximity (Arg128) to the DNA phosphate backbone, (**Fig. 5A**). In EMSAs, most of the pathological mutations in the β-hairpin (Lys113Glu, Arg115Trp, Arg116Gln, Arg116Pro, Lys125Glu), completely lose activity, in line with their DNA-binding roles highlighted in the cryo-EM structure. Instead, Arg128Gln partially loses its DNA-binding ability, and the Arg121Pro mutant binds DNA to the same extent as wild type DBD (**Fig. 7**). Arg128 is located at the end of the β-hairpin (**Fig. S1**), facing the DNA phosphate of T^-14^ at about 5.5 Å, thus its mutation may have an indirect negative effect on DNA-binding. Indeed, Arg128 is on one side, salt-bridged to the Asp130 side-chain and H-bonded to the carbonyl oxygen of Gly147, and is flanked by the sequence-specific Arg116 on the other side (**Fig. S9A**). Mutation to Gln is likely to alter the overall structure of the β-hairpin, preventing salt-bridge formation with Asp130 and possibly favoring H-bond interactions with Arg116, thus disturbing the latter residues in its key interactions with the palindromic NF-I site, as suggested by a significant reduction of DNA-binding shown in EMSA (**Fig. 7**). Arg121 is localized at a kink of the β-hairpin and interacts via a salt-bridge with OP1 T(8) in the DNA-bound form (**Fig. 5A**). The presence of a Pro residue in this position is compatible with preserving the kink in the protein backbone and the interaction with DNA may be compensated by the presence of a nearby Arg74 (**Fig. S9B**). Therefore, the pathological mechanism, at a cellular level, of this mutation, is likely to be more complex than a direct DNA-binding impairment (**Fig. 7**).

**Fig. 7.**
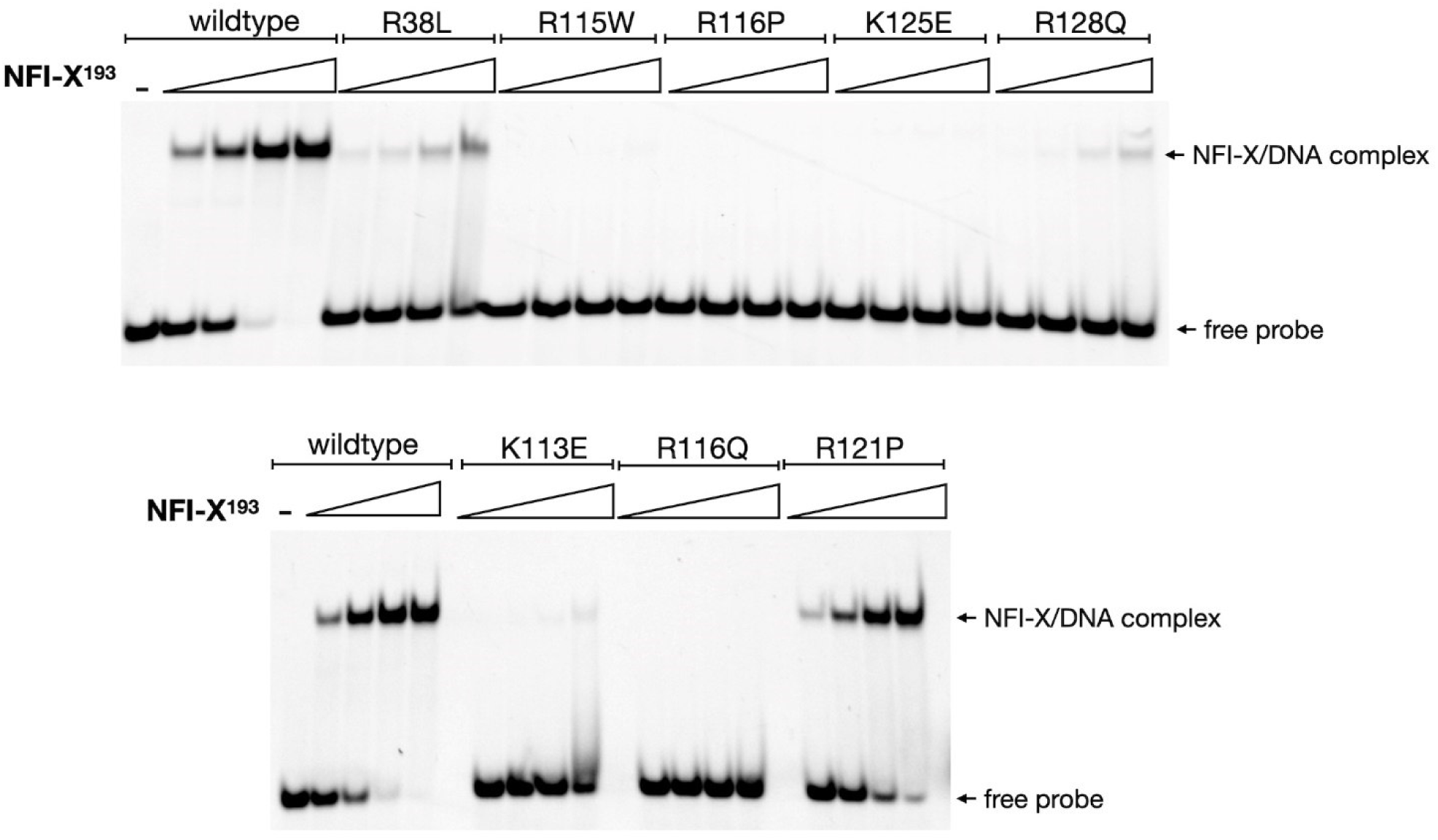
The effect of eight MALAN mutations on NFI-X^193^ binding to DNA. EMSA experiments were carried out for eight (Arg38Leu, Lys113Glu, Arg115Trp, Arg116Gln, Arg116Pro, Arg121Pro, Lys125Glu and Arg128Gln) NFI-X DBD mutations, identified in ten Malan syndrome individuals; each mutation is found in a different patient, except for Arg115Trp and Arg116Gln that were identified in two individuals each [6]. The effect of Malan mutations (single letter code) on DNA binding is shown in comparison with the NFI-X^193^ control, using 20 nM Cy5-labeled 31bp DNA probe (sequence shown in Figure 1) and 1X, 2X, 4X, and 8X concentrations of protein with respect to the probe. Bands corresponding to the free probe or the NFI-X^193^/DNA complex are indicated.

The antenna domain mutant (Arg38Leu) retains its DNA-binding ability, albeit with reduced affinity (**Fig. 7**). This is in line with the proposed mechanism of activation (**Fig. 6B**), whereby the Arg38Leu mutation would prevent the formation of the salt bridge with Asp124 in the DNA-unbound state, leading to a constantly activated TF, but the missing interaction between Leu38 and phosphate backbone in the DNA-bound state would then result in decreased DNA-binding affinity.

## Discussion

The NFI proteins shares little sequence homology with TFs for which structures have been determined, making the absence of structural information on this family particularly intriguing. Numerous studies have highlighted the pivotal role of NFIs in both development and disease, yet the mechanistic details of their ability to bind DNA in their target promoters has remained elusive until now. Although, some notions such as the requirement for protein dimerization and Zn^2+^ ions for DNA binding, have been deduced from EMSA-based studies, we can now provide the reasons, at a molecular level.

Here, we combined X-ray crystallography, cryo-EM, biophysical and mutational approaches to elucidate, for the very first time, the molecular mechanisms of DNA recognition by NFI-X at its palindromic NFI-site. The NFI-X DBD has a novel fold, based on a four helical bundle core domain, reminiscent of the Smad MH1 domain, and a α-loop-α antenna domain, specific for NFIs, inserted at the N-terminus of the DBD (**Fig. 1 and Fig. 4**). The core domain represents a stable molecular platform from which the antenna domain and a β-hairpin region protrude, forming a positively charged cleft deputed for DNA-binding (**Fig. S8**). In line with observations made for Smad MHI domains, NFI-X DBD contains a bound Zn^2+^ ion, coordinated by the CCCH motif (**Fig. 2C**), conserved within the NFI family (**Fig. 2A and Fig. 2D**). As for Smad proteins, this ion plays a stabilizing, structural role [41] and, accordingly its removal from NFI-X^193^ by chelation leads to protein instability and precipitation (results not shown). In particular, the Zn^2+^-binding site compacts the MH1 domain, properly orients a long loop that hosts the DNA-recognition β-hairpin towards the DNA major groove (**Fig. 2B**), and it stabilizes the C-terminal region containing the dimerization helix α5 (**Fig. 4A**). Our observations, finally rationalize previous biochemical and mutational data on human NFI-C that show that the three Cys coordinating residues are essential for DNA binding [36].

NFI-X binds to the NFI consensus sequence as a dimer, with each subunit inserting its β-hairpin into the major groove from the same side of the DNA (**Fig. 4A**). We showed that NFI-X DBD is a monomer in solution and dimerizes only upon DNA-binding (**Fig. 1C** and **Fig. S2**), via a mechanism that involves the swapping of the C-terminal helix α5 from each monomer (**Fig. 4B and Fig. 4C**). Removal of this helix reduces DNA-binding affinity by ten-fold (**Fig. 1A**) thus helix α5 appears to be both necessary and sufficient for NFI dimerization on DNA, but it is not sufficient for dimerization in solution (**Fig. S2**), although we cannot exclude the involvement of successive C-terminal residues for further dimer stabilization in the full-length protein. AlphaFold modelling of full-length NFI-X suggests that the regions following the dimerization helix α5 are intrinsically disordered (**Fig. S10**). In fact, previous coimmunoprecipitation data indicate that full-length NFI-C can dimerize with truncated mutant(s) devoid of DNA-binding ability [20]. It is, therefore, unsurprising that proteins derived from different NFI genes have been demonstrated to form heterodimers *in vitro* [25]. In this context, our structure suggests that the molecular mechanism of subunit selection for heterodimer formation must involve regions outside the DBD, where the NFI sequences diverge. The real occurrence of a NFI heterodimer in a cellular context is however still debated, although NFI genes have been reported to be simultaneously expressed in certain cell lines [25].

Our data reveal the unique activation mechanism and DNA-recognition mode exploited by NFI-X, based on the insertion of a β-hairpin loop contributed from each protein subunit of the dimer into the half site of the palindromic NFI-site, located in the DNA major groove (**Fig. 4A**). Base-reading interactions only involve two NFI-X residues (Arg116 and Lys 125) housed in the β-hairpin that contact the three G bases of each half site, rationalizing DNA consensus sequence mutations that confirm these bases as the most relevant for modulating DNA binding affinity [26]. Additionally, the methyl groups of the TT motif are hydrophobically stabilized by the facing Ala123. The DNA backbone is further stabilized by several ionic interactions involving residues belonging to both the NFI-X core (including the β-hairpin) and the antenna domains (**Fig. 5**). Our structural and mutagenesis data indicate these residues as functionally relevant, providing a rational interpretation, associated mostly to DNA-binding deficiency, for several pathological mutations identified in Malan Syndrome individuals (**Fig. 7**) [6,8,42].

Antenna domain residue Arg38, in particular, triggers a conformational change required to insert the β-hairpin loop into the major groove of the DNA. In unbound NFI-X, this residue participates in an intramolecular salt-bridge with Asp124, which is substituted by a phosphate group in the DNA-bound state. This switch, in the Arg38 binding state, unlocks the β-hairpin loop (containing Asp124) from the antenna domain and allows the Lys125 side-chain to relocate into the major groove to read the DNA consensus motif, together with Arg116. This ingenious activation mechanism is the only appreciable conformational change occurring in the unbound and DNA-bound states of NFI-X^193^. The structural dynamics of the β-hairpin loop required for DNA-binding is compensated by its anchoring to the core domain through Cys119, which is nestled in a hydrophobic pocked (**Fig. S1, and Fig. S9B)**. This result is in keeping with previous biochemical data showing that oxidation of this Cys residue (known as Cys-3) leads to reversible inactivation of the protein [22,27,28], and suggests that the discrepancy in the polarity of the oxidized residue and the surrounding environment can disrupt the structure of the β-hairpin loop.

The architecture of the dimeric NFI-X^193^/DNA complex also provides a structural explanation for the strong preference of a 5bp spacer in the NFI binding site (**Fig. 1C and Fig. 4A**). In fact, by EMSA, we observed that a 4bp spacer led to steric hindrance between the monomers and only one monomer was able to bind to a single half site (**Fig. 1C**). The reduced distance between the protomers would impede the swapping interaction and induce a clash between the C-terminal helix of one protomer with the antenna domain of the other (**Fig. S11**). In contrast, a spacer longer than 5bp would distance the C-terminal DBD helices from one another, preventing dimerization and thus decreasing DNA-binding affinity, as previously reported [21].

A final, interesting issue relates to the functional relevance of the super-helical structure present on the cryo-EM grids, whose formation was essential to solve the structure of such a small macromolecular complex (∼70 kDa) (**Fig. 3 and Fig. S7**). The stabilizing interactions of the super-helix involve base stacking at the 3’-and 5’-termini of adjacent DNA molecules, responsible for elongating the fibre, and protein-protein interactions, which bridge the two opposing DNA helices of the fibre. The latter interaction involves residues belonging to α1, α3, and the Zn^2+^-binding site of one subunit, with those of another subunit, a region opposite to that deputed to DNA-binding (**Fig. 3C**). Although, to our knowledge, no evidence has been reported (yet) for the presence of this supercomplex in a cellular context, this protein assembly found in our cryo-EM structure raises the striking possibility that NFI TFs mediate DNA-looping by packing, via protein-protein interactions, distant DNA regions that house NFI-sites. Interestingly, the cryo-EM fibre identified a hot-spot for protein-protein interactions that coincides with the region (α3) proposed to be involved in NFI-mediated activation of viral DNA replication through direct interaction of NFI with the viral preinitiation DNA polymerase complex [20]. In the NFI-X^193^/DNA complex, this region is solvent-accessible and is not blocked by the presence of bound-DNA. This is relevant since it provides the first structural evidence to support the proposed role of NFI as a DNA polymerase complex recruiter after binding at the origin of replication.

In conclusion, our data represent a Rosetta Stone, which not only structurally decodes a plethora of biochemical and functional data regarding NFI proteins but, in perspective, provides the basis for analysing new functional roles of NFIs and for the rational design of compounds that modulate the DNA-binding properties of this class of TFs, with possible applications in muscular dystrophy treatment in the case of NFI-X, and in related pathologies such as cancer, neurodevelopmental disorders, for other NFIs [43].

## Methods

### Protein Production of NFI-X DBD constructs

The cDNA encoding for the DBD of mouse NFI-X isoform 2 (UniProt code P70257), extending over amino acids 14 to 176 (NFI-X^176^), was chemically synthesized (Eurofins Genomics Srl.) and subcloned in-frame with a N-terminal maltose-binding protein (MBP) fusion tag, in the pMAL-cRI bacterial expression vector (New England Biolabs), via *EcoRI* and *HindIII* restriction sites. The codon-optimized, cDNA encoding for NFI-X^193^, covering residues 14 to 193, was synthesized and sub-cloned into the pET21-b (Novagen) via *NdeI* and *SalI* restriction sites, in frame with a N-terminal hexahistidine tag and Small ubiquitin-related modifier 3 tag (His-SUMO) (Genewiz, Azenta Life Sciences). Malan NFI-X^193^ mutants were generated by site-directed mutagenesis using Phusion DNA polymerase (Thermo Fisher Scientific), according to standard protocols, using the NFIX^193^ plasmid as a template.

Both NFI-X constructs were overexpressed in BL21 Shuffle T7 *E. coli* cells (New England Biolabs) grown at 30 °C in 1X (NFI-X^176^) or 2X (NFI-X^193^) Luria Bertani broth, supplemented with 100 μg/mL ampicillin. Protein expression was induced at optical densities of 0.6-0.8 at 600 nm wavelength, upon addition of 0.25 mM (NFI-X^193^) or 0.5 mM (NFI-X^176^) isopropyl β-d-1-thiogalactopyranoside (IPTG; Genespin Srl.) to cultures, pre-cooled to the 20 °C induction temperature, and further incubated overnight with shaking at 220 rpm.

Bacterial cells from 2L NFI-X^176^ cultures were harvested by centrifugation and lysed by sonication. NFI-X^176^ was purified from the supernatant using a 5 mL HiTrap Heparin HP affinity column (Cytiva), pre-equilibrated in Binding Buffer A (50 mM HEPES pH 7.0, 200 mM NaCl and 2 mM dithiothreitol (DTT) using an AKTA pure™ FPLC system (Cytiva). The protein was eluted using a linear salt gradient, achieved by mixing Buffer A with Buffer B (50 mM HEPES pH 7.0, 1 M NaCl and 2 mM DTT). Fractions containing NFI-X^176^ were pooled and incubated with 50 U thrombin protease (Sigma Aldrich), with gentle rotation overnight at 20 °C. The cleavage reaction was diluted with 50 mM HEPES pH 7.0 and 2 mM DTT to lower the NaCl concentration to 200 mM. Cleaved NFI-X^176^ was separated from the MBP fusion tag and intact MBP fused constructs by a second Heparin affinity chromatography step, using analogous, forementioned purification conditions. Fractions containing pure NFI-X^176^ were pooled, concentrated to 500 μl with an Amicon ultracentrifugal concentrator (Merck Millipore) with a MW cut-off of 10000, and further purified by SEC on a Superdex 200 Increase 10/300 column (Cytiva), pre-equilibrated in Buffer A.

Bacterial cells from 2L NFI-X^193^ cultures were mechanically lysed at 25 mPa using a cell disruptor (Constant Systems Ltd) and sonication. NFI-X^193^ was purified from the clarified supernatant using a 1 mL Histrap HF column (Cytiva), pre-equilibrated in Buffer A (20 mM sodium phosphate pH 7.4, 20 mM imidazole, 0.5 M NaCl, 1 mM DTT). Elution was carried out over a linear gradient, performed mixing Buffer A with Buffer B (Buffer A containing 0.5 M imidazole). Peak fractions were pooled and dialyzed overnight into SUMO protease cleavage buffer (50 mM Tris pH 7.4, 200 mM NaCl, and 1 mM DTT). Removal of the His-SUMO tag was achieved upon incubation of the fusion protein at room temperature for 4h with in-house, recombinantly produced *Saccharomyces cerevisiae* SUMO protease at a protease:protein molar ratio of 1:200, respectively. Cleaved NFI-X^193^ was separated from the His-SUMO fusion tag via affinity chromatography on a 5 mL Heparin HP column (Cytiva), using the conditions used to purify NFI-X^176^. Fractions containing purified NFI-X^193^ were concentrated to 500 μl and further purified on a ProteoSEC 6-600 HR 10/300 column (Neobiotech), pre-equilibrated in 50 mM Tris pH 7.4, 200 mM NaCl and 1 mM DTT. Molecular weights were calculated from calibration curves generated using the Bio-rad Gel Filtration Standards (Bio-rad Laboratories), using standard protocols. Malan mutants were expressed and purified as for NFI-X^193^.

### NFI-X/DNA complex assembly for crystallization and cryo-EM

Co-crystallization experiments of NFI-X^176^ in complex with *NFI-31bp* were unsuccessful, therefore a shorter 23bp DNA fragment (*NFI-23bp*) with sticky ends (forward strand: 5’-GTCTCT<u>TTGGC</u>AGGCA<u>GCCAA</u>CC-3’; reverse strand 5’-ACGG<u>TTGGC</u>TGCCT<u>GCCAA</u>AGAG-3’), containing the two NFI half sites (underlined) were set up. 23 bp is long enough to be bound by NFI-X, but short enough not to interfere with crystallogenesis (**Fig. S3**). Sticky ends were selected to stabilize and promote crystal lattice formation. The protein DNA complex was assembled, mixing protein:DNA at a 2:1 molar ratio and by progressively decreasing the buffer salt concentration to 50 mM, to promote complex assembly by repetitive concentration and dilution cycles with buffer, before final concentration to 15 mg/mL.

For Cryo-EM, the NFI-X^193^ complex was assembled with the 31bp DNA sequence used in EMSAs (**Fig. 1**). NFI-X^193^ fractions, derived from the final Heparin affinity chromatography step, were mixed with DNA at a molar ratio of 2:1. The high salt present in the Heparin chromatography elution Buffer B was gradually lowered to promote complex formation, by stepwise dialysis into 50 mM Tris pH 7.4, containing 1 mM DTT and progressively lower NaCl concentrations (300 mM, 150 mM and 75 mM NaCl). Each dialysis step was carried out at 4 °C in 2L dialysis buffer using a Spectra-Por® dialysis membrane (Spectrum™) with a 3.5 kDa cut-off and buffer exchanges every 2h, with the last step overnight. The dialysate was concentrated to 500 μl and loaded onto ProteoSEC 6-600 HR 10/300 column (Neobiotech), pre-equilibrated in 20 mM Tris pH 7.4, 150 mM NaCl and 1 mM DTT. NFI-X^193^-DNA complex-containing fractions were concentrated to 10 mg/mL.

### Circular dichroism (CD)-based thermal denaturation studies

The thermostability of protein unfolding was investigated by monitoring ellipticity variations at a single wavelength of 222 nm over a temperature gradient of 20 °C to 95 °C (1 °C/minute ramp rate). Unfolding curves are represented as change in ellipticity versus temperature; 0.2 mg/mL protein samples (20 mM Tris pH 7.4, 200 mM NaCl and 1 mM DTT) were placed in 1 cm quartz cuvettes and CD spectra were recorded using a Jasco J-810 instrument equipped with a PFD-425S Peltier temperature controller (Jasco).

### Electrophoretic Mobility Shift Assays

DNA-binding reactions were assembled on ice using DNA probes carrying the NFI consensus sequence. The probe was designed and optimized, based on the sequence of the promoter region of NFI-X isoforms 3 and 4 [51]. Starting from this sequence (5′-TGGGTCTCT**<u>TTGGC</u>**AGCA**<u>GCCAC</u>**CCAGCAAA-3), 1 nucleotide (guanine) insertion was made after adenine 18 to lengthen the spacer region to 5 nt and another substitution was made (c23a) to make the two half sites palindromic (in bold font and underlined), based on previous studies correlating DNA sequence to NFI binding affinity [21,52]. Probes were labelled with Cyanine 5 (Cy5) at the 5’-end of the forward strand and annealed with the unlabeled reverse strands, using standard protocols. Dose response experiments were performed by combining 14 μL of a premixed Binding Mix (BM), containing 20 nM NFI Cy5-probe, 20 mM Tris-HCl pH 7.5, 50 mM NaCl, 5 mM MgCl2, 30 mM KCl, 0.5 mM EDTA, 6.5% *(v/v)* glycerol, 2.5 mM DTT, 0.1 mg/mL BSA, 10 ng Poly(dIdC) (Thermo Fisher Scientific), with 2 μL purified NFI-X^176^ or NFI-X^193^ at varying final concentrations (0 nM, 60 nM, 180 nM, 540 nM, 1.6 μM). Reactions were incubated at 30 °C for 30 min in the dark and electrophoresis was performed in the dark on a 6% Native gel run at 100V in 0.25X Tris/Borate/EDTA buffer. Cy5 fluorescence was detected using a Chemidoc MP imaging apparatus (Bio-Rad Laboratories) and band intensity was quantified with ImageLab software.

Competition EMSAs were carried out, as described above, in 14 μl binding mix containing 20 nM NFI-X^193^ and 20 nM Cy5-labelled 31bp DNA, incubated with 2 μl unlabelled 31bp competitor or 31bp random probe (**Fig. 1B**) at increasing concentrations (6.25X, 12.5X, 25X and 50X with respect to the labelled probe).

### Flame atomic absorption spectroscopy (FAAS)

The presence of Zn^2+^ was assessed in purified NFI-X^176^ by calibrated FAAS. A homogeneous solution of 0.2 mg/mL NFI-X^176^ was exposed to wet decomposition. To the protein sample, 5 mL of nitric acid (65% *(v/v))* were added and the resulting solution was gently boiled on a hot plate until the red fumes coming from the beaker terminated (1 h). After cooling, 5 mL freshly prepared mixture of 65% nitric acid and 37% hydrochloric acid (1:3) were added and the solution was warmed on a hot plate for 2 h. After cooling, about 3 mL of hydrogen peroxide (30%) were added and the solution was boiled again to evaporate until a small portion remained. After cooling, the resulting clear digested solution was quantitatively diluted to a final volume of 10 mL with 2% nitric acid before being analysed in duplicate by FAAS. Nitric acid, hydrochloric acid and hydrogen peroxide were purchased from Sigma Aldrich.

The calibration curve was prepared using five standards (including the blank). Working calibration solutions were freshly prepared using appropriate stepwise dilutions of standard Zn stock solution (Ultra grade, 1000 mg/L, 2% HNO_3_, Perkin-Elmer). Working standards were as follows: 0.25, 0.5, 1 and 1.4 ppm diluted in 2% *(v/v)* nitric acid.

Data were collected by using an atomic absorption spectrometer (PinAAcle 900T, Perkin-Elmer) equipped with deuterium lamp for background correction, air-acetylene flame and zinc hollow-cathode lamp operating at 213.9 nm. The linear range was 0.01-2 mg/L. The measurements were performed in triplicate, and the mean was automatically calculated using the standard calibration graph.

### Crystallization and 3D Structure Determination of NFI-X^176^

Sitting-drop vapor-diffusion experiments were set up using an Orxy8 robot (Douglas Instruments) purified NFI-X^176^/DNA (*NFI-23bp*) complex (15 mg/mL). Crystals grew at 4 °C in an optimized condition based on condition B2 of the JCSG Screen (Molecular Dimensions): 0.2 M sodium thiocyanate (NaSCN), 0.1 M HEPES pH 7.0, 22% *(w/v)* PEG3350 in 96-well, flat-bottomed microbatch plates (Greiner-BIO-ONE) at a protein:precipitant ratio of 50% *(v/v)* and 9 mL of a 1:1 paraffin:silicon oil mixture as sealing agent. Crystals were soaked in cryoprotectant solution (0.2 M NaSCN, 0.1 M HEPES pH 7.0, 22% *(v/v)* PEG 3350, 20 % *(v/v)* glycerol) and flash frozen in liquid nitrogen.

X-ray diffraction data collections were collected at the XRD2 beamline of the ELETTRA Synchrotron, Trieste (Italy), equipped with a Pilatus 6M hybrid pixel area detector (Dectris). Data were indexed, integrated, and scaled using XDS [53]. The statistics for the data collection and data reduction are summarized in **Table S1**.

The only NFI-X DBD structural homolog is the MH1 domain of Smad proteins, that shares low (<20%) sequence identity, and attempts to phase the NFI-X data with a Smad model by Molecular Replacement failed. The structure was then solved by SAD phasing in the *P*12_1_1 space group, exploiting the anomalous signal of bound zinc atoms at a wavelength of 1.2705Å. The heavy atom substructure was determined using SHELX C/D/E [54,55]. ARP/wARP was used for automated model building [56]. The structure was refined using REFMAC5, Phenix.refine and multiple rounds of manual model building in Coot [57–59] (Table S1). MolProbity and PISA were used to assess the stereochemical quality and protein quaternary assembly, respectively [39,60]. Atomic coordinates and structure factors were deposited under PDB entry 7QQD.

### Cryo-EM sample preparation

3-{Dimethyl[3-(3α,7α,12α-trihydroxy-5β-cholan-24-amido)propyl]azaniumyl}propane-1-sulfonate (CHAPS) detergent was added at its critical micelle concentration of 8 mM to 4 mg/mL purified NFI-X^193^/DNA sample. Quantum foil R 0.6/1 Cu 300 grids were glow-discharge treated for 30 s at 30 mA, using the GloQube system (Quorum Technologies). Approximately, 3 mL sample were applied onto several grids, under controlled conditions of 4 °C and 100% relative humidity. Various blotting times were tested to optimize sample distribution and minimize excess liquid. Subsequently, prepared grids were rapidly vitrified in liquid ethane utilizing a Vitrobot Mark IV (Thermo Fisher Scientific). Finally, vitrified grids were transferred and stored in liquid nitrogen until further analysis.

### Cryo-EM data collection and image processing

Cryo-EM data were acquired from prepared grids using an in-house FEI Talos Arctica 200 kV FEG transmission electron microscope (Thermo Fisher Scientific). An electron beam, operating at 200 kV and a calculated electron dose of 40 electrons per square angstrom (e^-^/Å^2^), was directed through the sample and reached a direct-electron detector for image capture. Data acquisition was carried out using EPU-2.8 automated data collection software (Thermo Fisher Scientific) and a series of movies, each composed of 40 individual frames, were recorded. The microscope was set to a nominal magnification of 120,000x, which corresponded to a pixel size of 0.889 angstroms per pixel at the specimen level. During data collection, defocus values were systematically adjusted within the range of-0.8 to-2.2 μm. This range of defocus values was selected to optimize image contrast and resolution, allowing for the capture of structural details within the protein-DNA complex.

Micrographs were initially selected based on visual quality inspection. Movie drift correction and CTF parameters determination were performed in Relion 3.1 [61]. The fibrils were manually picked and extracted with an inter-box distance of 44 Å along the helical axis into overlapping boxes of 400×400 pixels, resulting in 201,833 extracted segments. Such particles were then imported in CryoSPARC for 2D classification and 3D reconstruction CRY [62]. 2D class averages were used to discard bad quality particles, resulting in 172,304 selected segments, that were used to perform an *ab-initio* reconstruction. The *ab-initio* volume was used as an initial model to determine the helical symmetry parameters by using the *Symmetry search* tool in CryoSPARC, revealing that the helix was right-handed, with a helical rise of 44.51 Å, a twist of 142.5°, 2.5 subunits per turn and a pitch of 112.4 Å **(Fig. S7)**. These values were given as inputs for the *Helix Refine* tool and using the *ab-initio* model as input volume, that generated a first map at 4.2 Å resolution. From inspecting this volume, we observed the presence of a symmetry binary axis, orthogonal to the helix axis. Therefore, helical refinement was repeated imposing D1 symmetry, resulting in a final map at 3.86 Å resolution, with optimized helical twist and rise values of 142.672° and 44.618 Å, respectively.

### Cryo-EM Model building and Refinement

Two copies of the crystallographic structure of NFI-X^176^ (PDB: 7QQD) were initially fitted into the EM density of a single NFI-X^193^ dimer. Additional C-terminal residues (175-181), that were not present in the NFI-X^176^ construct and/or crystal structure, were manually built in COOT [57], together with a linear DNA molecule containing the NFI-X consensus sequence. The 5’ and 3’ DNA moieties were adjusted in COOT since they slightly deviated from linearity. The model of the complex was then iteratively rebuilt and all-atom real-space refined using stereochemical and NCS restraints within PHENIX [59]. The model of the dimer was then copied and fitted into the unmodelled EM density to obtain an atomic model of the fibril. The resolution was improved with an additional helical refinement imposing D1 symmetry, obtaining a final volume of NFI-X^193^-DNA complex fibrils at 3.86 Å resolution.

Cryo-EM data collection and processing statistics are summarized in **Table S2**. The PDB and EMDB accession codes of the NFI-X^193^/DNA complex are PDB ID 9QKY, EMD-53223, respectively.

## Abbreviations

CD: circular dichroism
DBD: DNA-binding domain
EMSA: electromobility shift assay
FAAS: Flame Atomic Absorption Spectrometry
MALNS: Malan syndrome
MSS: Marshall-Smith Syndrome
MW: molecular weight
NFI: Nuclear factor 1
NFI-X: Nuclear factor 1 X
SEC: Size exclusion chromatography
TAD: transactivation domain
TF: transcription factor
T_M_: melting temperature.

## Acknowledgements

We thank the Trieste staff on beamline XRD2. Cryo-EM studies were carried out using the Unitech NOLIMITS facility at the University of Milan. LJG is grateful for funding from “Linea 2” Università degli Studi di Milano.

## Notes

### Competing Interest Statement

The authors have declared no competing interest.

