## Supplemental figures for "Structural bases for Nuclear Factor 1-X activation and DNA recognition. Prototypic insight into the NFI transcription factor family"

**Table S1. X-ray data collection and refinement statistics.**

**Table S2: Cryo-EM data collection and processing.**

**Fig. S1. NFI sequence alignment.**

**Fig. S2: Biochemical and biophysical characterisation of NFI-X DBD constructs.**

**Fig. S3: Effect of DNA length on NFI-X$^{176}$ binding affinity.**

**Fig. S4. Zinc ions are bound to NFI-X$^{176}$.**

**Fig. S5. DNA recognition by Smad TF.**

**Fig. S6. Crystal packing of NFI-X$^{176}$.**

**Fig. S7. Cryo-EM processing workflow for the helical reconstruction of NFI-X$^{193}$-DNA fibril.**

**Fig. S8. Electrostatic surface representation of NFI-X$^{193}$ in complex with DNA.**

**Fig. S9. Structural position and interactions of Malan Syndrome residues Arg128 and Arg121.**

**Fig. S10. AlphaFold prediction of full-length NFI-X.**

**Fig. S11. Model of the NFI-X$^{193}$ dimer bound to NFI consensus sequence with a shorter, 4bp-spacer.**

**Table S1. X-ray data collection and refinement statistics.**

| | (PDB code 7QQD) |
|---|---|
| **Crystal** | |
| Space group | $P12_11$ |
| Unit cell dimensions<br>$a$, $b$, $c$ (Å); α, β, γ (°) | 43.2, 98,4, 61,6<br>90.0, 92.7, 90.0 |
| **Data collection** | |
| Beamline | ELETTRA XDR2 |
| Wavelength (Å) | 1.27 |
| Resolution (Å) | 2.70 (2.83-2.70) |
| Unique reflections | 14029 (1374) |
| Total reflections | 92576 |
| $^{\Psi}R_{merge}$ | 0.084 (0.868) |
| $^{\#}R_{meas}$ | 0.101 (1,035) |
| $I/\sigma(I)$ | 13.2 (2.0) |
| $^{+}CC_{1/2}$ | 0.968 (0.817) |
| Completeness (%) | 99.3 (98.9) |
| Average redundancy | 6.6 (6.6) |
| **SAD Phasing** | |
| Anom. completeness (%) | 91.8 (91.1) |
| $R_{anom}$ | 0.052 (0.543) |
| $CC_{anom}$ | 0.523 (0.061) |
| Average FOM<br>(after automated fitting of 150 Ala residues) | 0.623 |
| **Refinement** | |
| Resolution (Å) | 49.2-2.70 (2.80-2.7.0) |
| No. reflections | 14029 (1374) |
| $R$ / $R_{free}$ | 0.206/0.262 |
| No. molecules (non-H atoms) in the asymmetric unit | 2 |
| Protein | 2 |
| Water | 35 |
| Zinc ions | 4 |
| HEPES | 1 |
| Average B factors (Å$^2$) | |
| Protein | 76.1 |
| Water | 70.6 |
| Zinc ions | 80.0 |
| HEPES | 70.4 |
| RMSD from ideal values | |
| Bond lengths (Å) | 0.016 |
| Bond angles (°) | 1.96 |
| **Ramachandran** | |
| Favored (%) | 93.44 |
| Allowed (%) | 99.16 |
| Outliers (%) | 0.64 |

* Values in parenthesis correspond to the high-resolution shell. For cross-validation, 5% experimental reflections were randomly selected to calculate the $R_{free}$ value.
$^{\Psi}$ $R_{merge} = \Sigma_h \Sigma_i |<I_h> - I_{h,i}| / \Sigma_h \Sigma_i I_{h,i}$
$^{\#}$ $R_{meas} = \Sigma_h [N_h/(N_h - 1)]^{1/2} \Sigma_i|<I_h> - I_{h,i}|/\Sigma_h \Sigma_i I_{h,i}$, where $N_h$ is the data multiplicity.
$^{+}$ $CC_{1/2}$ is the correlation coefficient of the mean intensities between two random half-sets of data.

**Table S2: Cryo-EM data collection and processing.**

| | (PDB code 9QKY) |
|---|---|
| **Data collection and processing** | |
| Microscope | FEI Talos Arctica |
| Magnification | 120,000 |
| Voltage (kV) | 200 |
| Electron exposure ($e^-$/Å$^2$) | 40 |
| Nominal defocus range (μm) | 0.8 - 2.2 |
| Pixel size (Å) | 0.899 |
| Imposed Symmetry | D1 |
| Micrographs (no.) | 1225 |
| Initial segment images (no.) | 201,833 |
| Final segment images (no.) | 164,794 |
| Map resolution (Å) | 3.86 |
| FSC threshold | 0.143 |
| Map resolution range (Å) | 3.2 - 6.5 |
| **Refinement** | |
| Initial model used (PDB code) | 7QQD |
| Model resolution (Å) | 2.7 |
| **Model composition** | |
| Non-hydrogen atoms | 16276 |
| Protein residues | 1352 |
| Nucleotides | 248 |
| Ligands | 8 |
| ***B* factors (Å$^2$)** | |
| Protein | 70.93 |
| Nucleotide | 155.83 |
| Ligand | 92.40 |
| **R.m.s. deviations** | |
| Bond lengths (Å) | 0.003 |
| Angles (°) | 0.581 |
| **Validation** | |
| MolProbity score | 1.62 |
| Clashscore | 6.35 |
| Poor Rotamers (%) | 0.65 |
| **Ramachandran** | |
| Favored (%) | 96.11 |
| Allowed (%) | 3.89 |
| Outliers (%) | 0 |
| **Model vs Data** | |
| CC(mask) | 0.85 |
| CC(box) | 0.75 |
| CC(peaks) | 0.56 |
| CC(volume) | 0.81 |
| CC(main chain) | 0.83 |
| CC(side chain) | 0.81 |

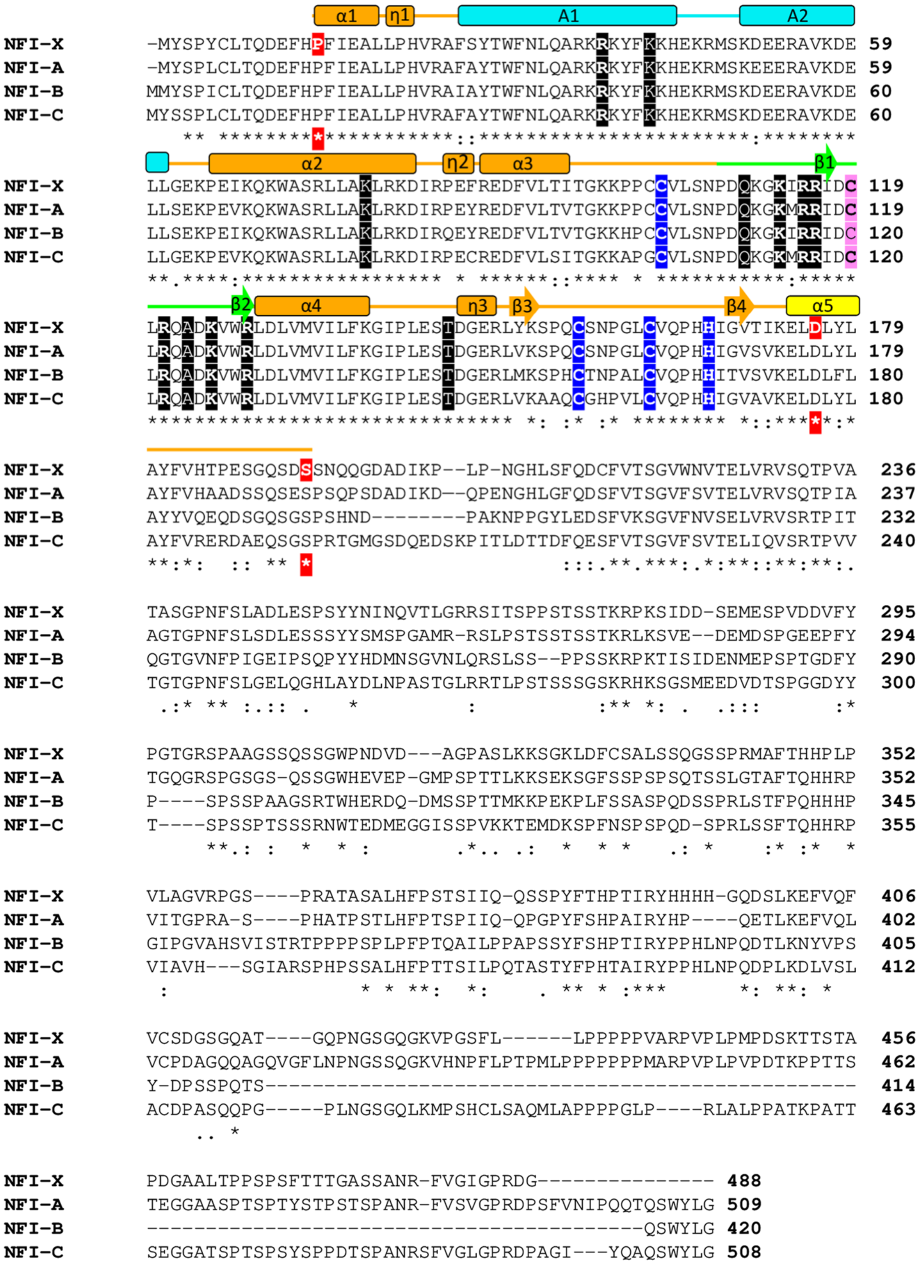


**Figure S1. NFI sequence alignment.** A multiple sequence alignment was carried out with mouse NFI-X (Uniprot code P70257), human NFI-A (Uniprot code Q12857), NFI-B (Uniprot code O00712) and NFI-C (Uniprot code P08651), using ClustalOmega [24]. Conserved residues are indicated by asterisks (identical) and colons/dots (similar). The first amino acid (Pro14) and the C-terminal residues of NFI-X$^{176}$ and NFI-X$^{193}$ constructs are indicated in white bold font with red shading. The three cysteine residues Cys103, Cys156 and Cys162 (corresponding to Cys-2, Cys-4 and Cys-5) and His167, all involved in Zinc ion coordination, are highlighted in white bold font with black shading. Cys119 (corresponding to Cys-3) is highlighted in bold font with pink shading. Amino acid residues that participate in DNA binding are in white and shaded in blue, and in bold font if their mutation is associated with Malan syndrome [6]. The secondary structure elements of NFI-X are indicated and labeled, in accordance with secondary structure designation with PDBsum [49]. The MH1 core domain is shown in orange, the antenna domain in cyan, the dimerization helix in yellow, and the β-hairpin in green.

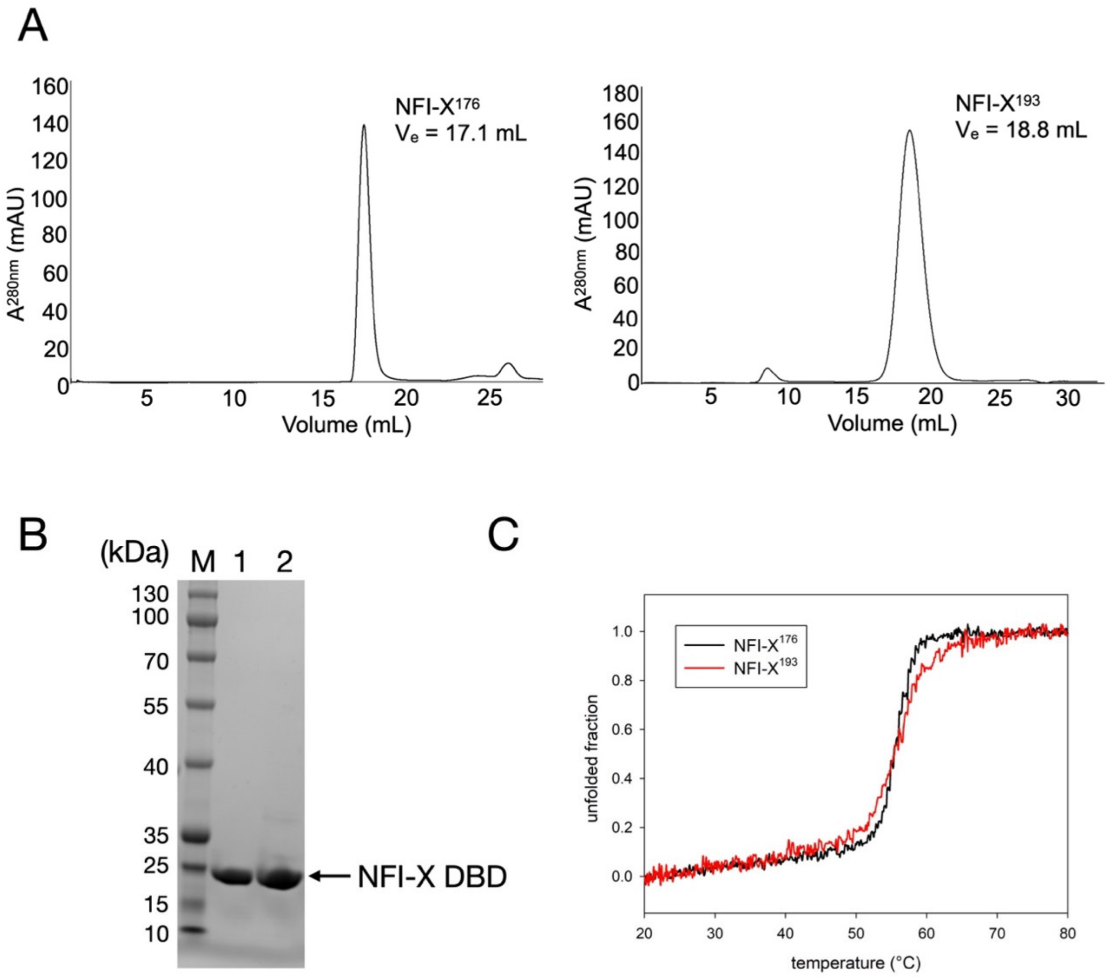


**Fig. S2. Biochemical and biophysical characterisation of NFI-X DBD constructs. A)** Size exclusion chromatography (SEC) profile of NFI-X$^{176}$ (left panel) and NFI-X$^{193}$ (right panel), using a Superdex 200 Increase 10/300 column (Cytiva) and a ProteoSEC 6-600 HR 10/300) column (Neo Biotech), respectively, as described in the Methods section. Elution profiles show the absorbance at 280 nm (mAU) as a function of volume (mL). NFIX$^{176}$ and NFIX$^{193}$ elute at elution volumes ($V_e$) of 17.1 mL and 18.8 mL, respectively, corresponding to a monomer in solution; **B)** SDS-PAGE analysis of purified NFI-X$^{176}$ (Lane 1) and NFI-X$^{193}$ (Lane 2) samples on a 4-20% Novex™ Tris-Glycine Mini Protein gel (Invitrogen). NFI-X$^{176}$ (Lane 1) and NFI-X$^{193}$ (Lane 2) migrate with a MW around 23 kDa, in line with their calculated MWs of 19.5 kDa and 21.4 kDa, respectively. The MWs for the PageRuler Prestained Protein Ladder (Invitrogen) are shown; **C)** The thermostability of NFI-X$^{176}$ (black line) and NFI-X$^{193}$ (red line) was deduced by circular dichroism, following the change in ellipticity variations at a single wavelength of 222 nm, corresponding to loss of α-helices, as a function of temperature. Unfolding curves are expressed as the proportion of total unfolded protein versus temperature (°C).

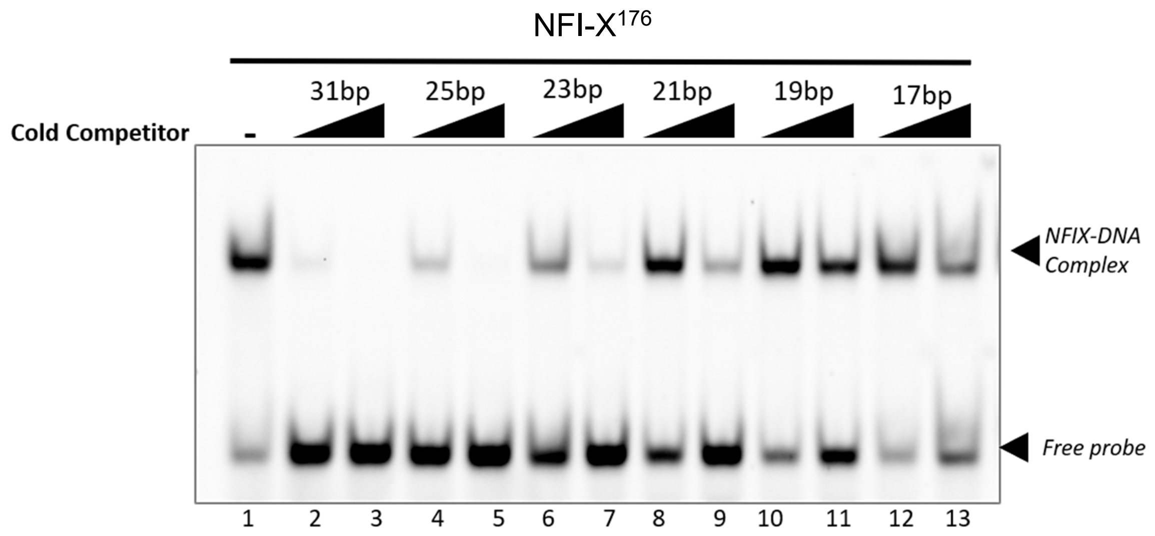


Cy5-labelled 31bp ds DNA probe:

31bp: 5'[Cy5]-GGGTCTCT**TTGGC**AGGCA**GCCAA**CCAGCAAA-3'

Unlabelled competitors:

31bp: 5'-GGGTCTCT**TTGGC**AGGCA**GCCAA**CCAGCAAA-3'

25bp: 5'-TCTCT**TTGGC**AGGCA**GCCAA**CCAGC-3'

23bp: 5'-CTCT**TTGGC**AGGCA**GCCAA**CCAG-3'

21bp: 5'-TCT**TTGGC**AGGCA**GCCAA**CCA-3'

19bp: 5'-CT**TTGGC**AGGCA**GCCAA**CC-3'

17bp: 5'-T**TTGGC**AGGCA**GCCAA**C-3'

**Fig. S3. Effect of dsDNA length on NFI-X$^{176}$ binding affinity.** EMSA competition experiments were carried out to assess NFI-X$^{176}$ binding affinity to varying lengths of dsDNA specific competitors (from 31 bp to 17 bp) for which the sequences are shown. All competitors contained the same palindromic NFI binding site (bold) with a 5 bp spacer (underlined). 20 nM NFI-X$^{176}$, mixed with 20 nM Cy5-labelled 31 bp DNA probe, was incubated in the presence of unlabelled DNA competitors at a 10- and 50-fold molar excess, with respect to the labelled probe, and run on a 6% polyacrylamide gel in 0.25X TBE. DNA binding was assessed as compared to the control sample, in the absence of competitor (lane 1).

A

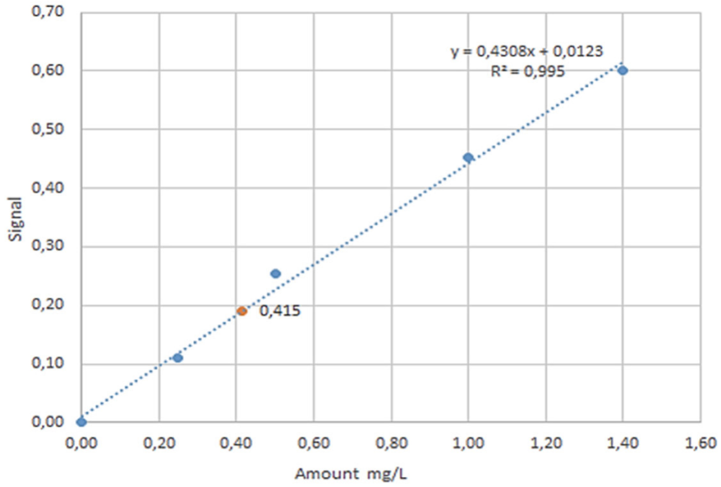


B

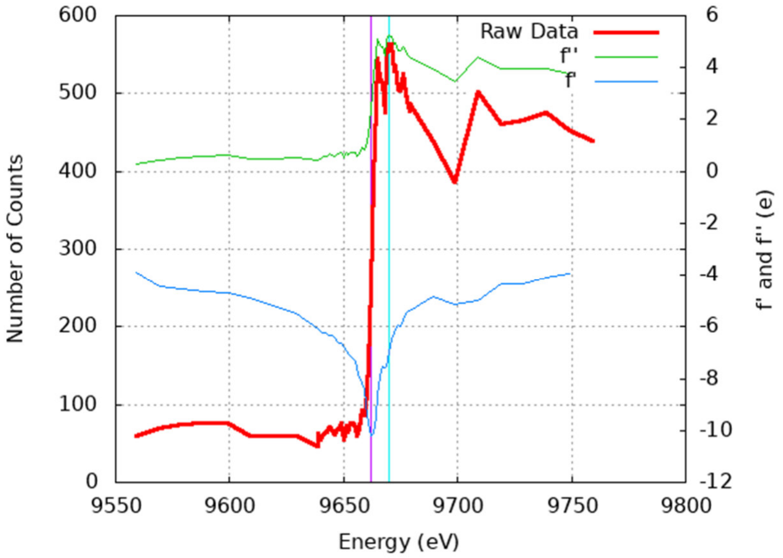


**Fig. S4. Zinc ions are bound to NFI-X[176]. (A)** Correlation coefficient of the zinc calibration curve by FAAS analysis. The amount of zinc in NFI-X[176] sample was estimated to be 0.415 mg/L (RSD: 4.7%). **(B)** XFR plot of the absorption spectrum of Zn K-edge. The graph plots the number of photons counts versus their energy (eV). Scattered electrons (e) submitted to derivative operators to calculate the spectra (f' and f''). Absorption at the zinc energy edge, 9670.0 eV, indicates that zinc is bound in the NFI-X[176] crystal.

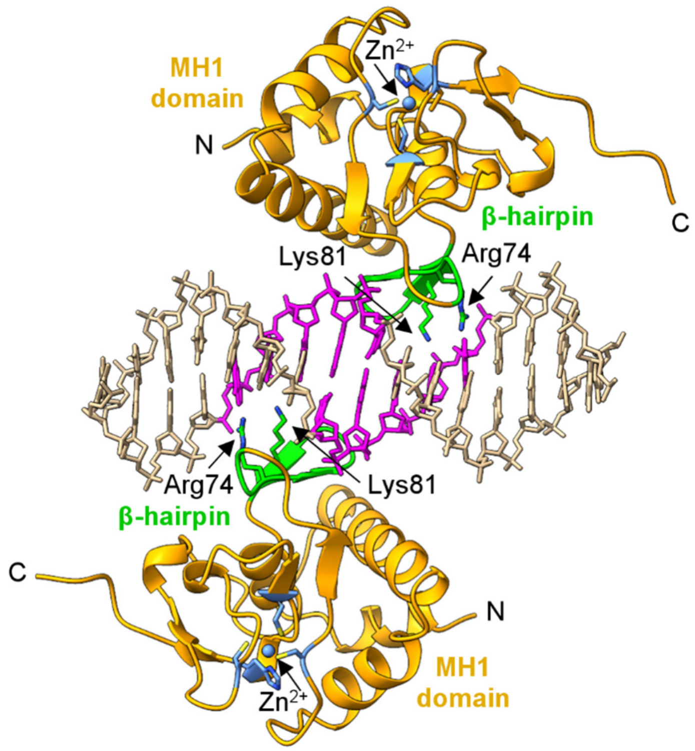


**Fig. S5. DNA recognition by Smad TF.** Ribbon representation of the crystal structure of human Smad3 dimer (PDB: 5OD6), bound to DNA [46]. Color code for secondary structure elements in the MH1 domain as in Fig. 2. The β-hairpin loop (green), involved in DNA-site recognition (magenta), is highlighted. Arg74 and Lys81, conserved in NFI-X[193] (Arg116 and Lys125), are shown as green sticks with nitrogen atoms in blue. The $Zn^{2+}$ ion is shown as a blue sphere and coordinating residues (CCCH motif) are shown as light blue sticks. The termini of protein are labelled. This figure was generated using ChimeraX version 1.8 [50].

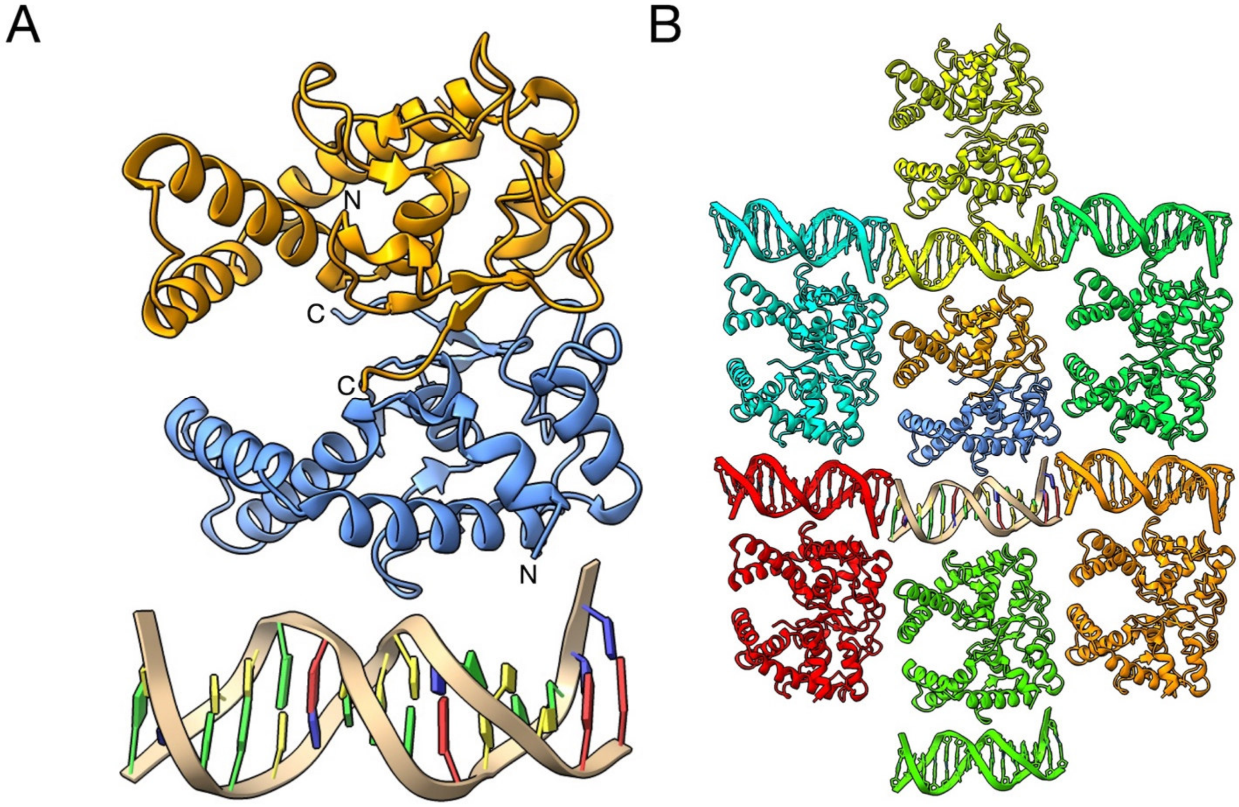


**Fig. S6. Crystal packing of NFI-X^176^. (A)** The crystal asymmetric unit (ASU) of the $P2_1$ crystal of NFI-X$^{176}$ (PDB: 7QQD), showing two monomers (in blue and orange ribbons) and the unbound, 15bp dsDNA fragment (ribbon representation) **(B)** Illustration of crystal lattice packing in the monoclinic $P2_1$ form, underlining the head to tail organization of the dsDNA packed at the crystal contacts. Each asymmetric unit is shown in a different color, with the ASU shown in panel A in the centre. This figure was generated using ChimeraX version 1.8) [50].

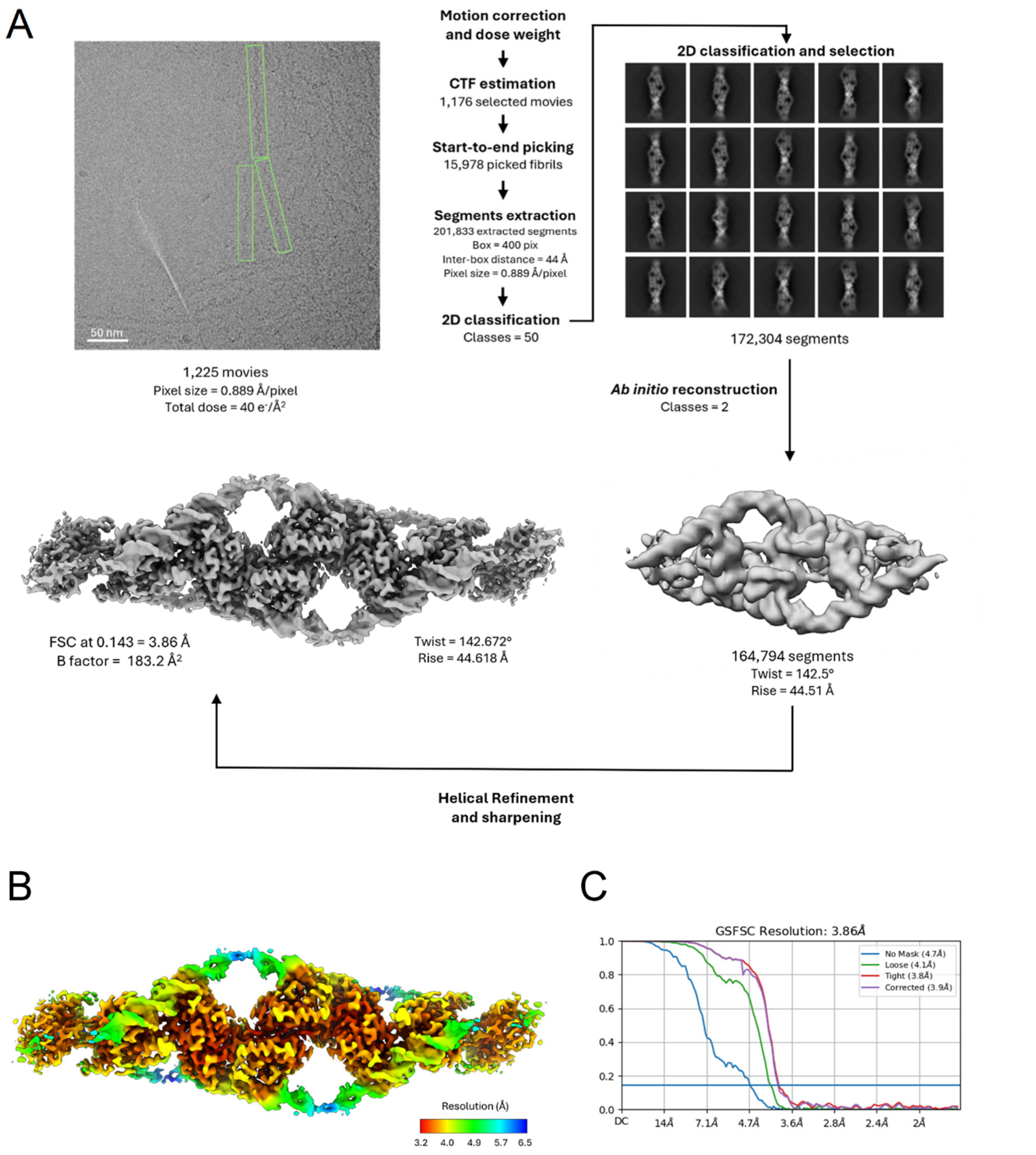


**Fig. S7. Cryo-EM processing workflow for the helical reconstruction of the NFI-X$^{193}$-DNA fibril. (A)** Schematic representation of the key steps and parameters in the cryo-EM data processing workflow for helical reconstruction. The figure includes a representative micrograph (scale bar: 50 nm), selected 2D class averages, *ab initio* volume, used as an initial model and for determining the helical symmetry parameters (twist = 142.5°; rise = 44.51 Å), and the final sharpened map obtained through helical refinement, which yielded optimized symmetry parameters (twist = 142.672°; rise = 44.618 Å). All steps were performed in CryoSPARC [62]; **(B)** The final reconstruction, colored according to local resolution, ranging from 3.2 to 6.5 Å; **(C)** The gold-standard Fourier shell correlation (GSFSC) curve of the final helical reconstruction, indicating a resolution of

3.86 Å at a Fourier shell correlation (FSC) threshold of 0.143. Cryo maps in Panels A and B were prepared using ChimeraX version 1.8) [50].

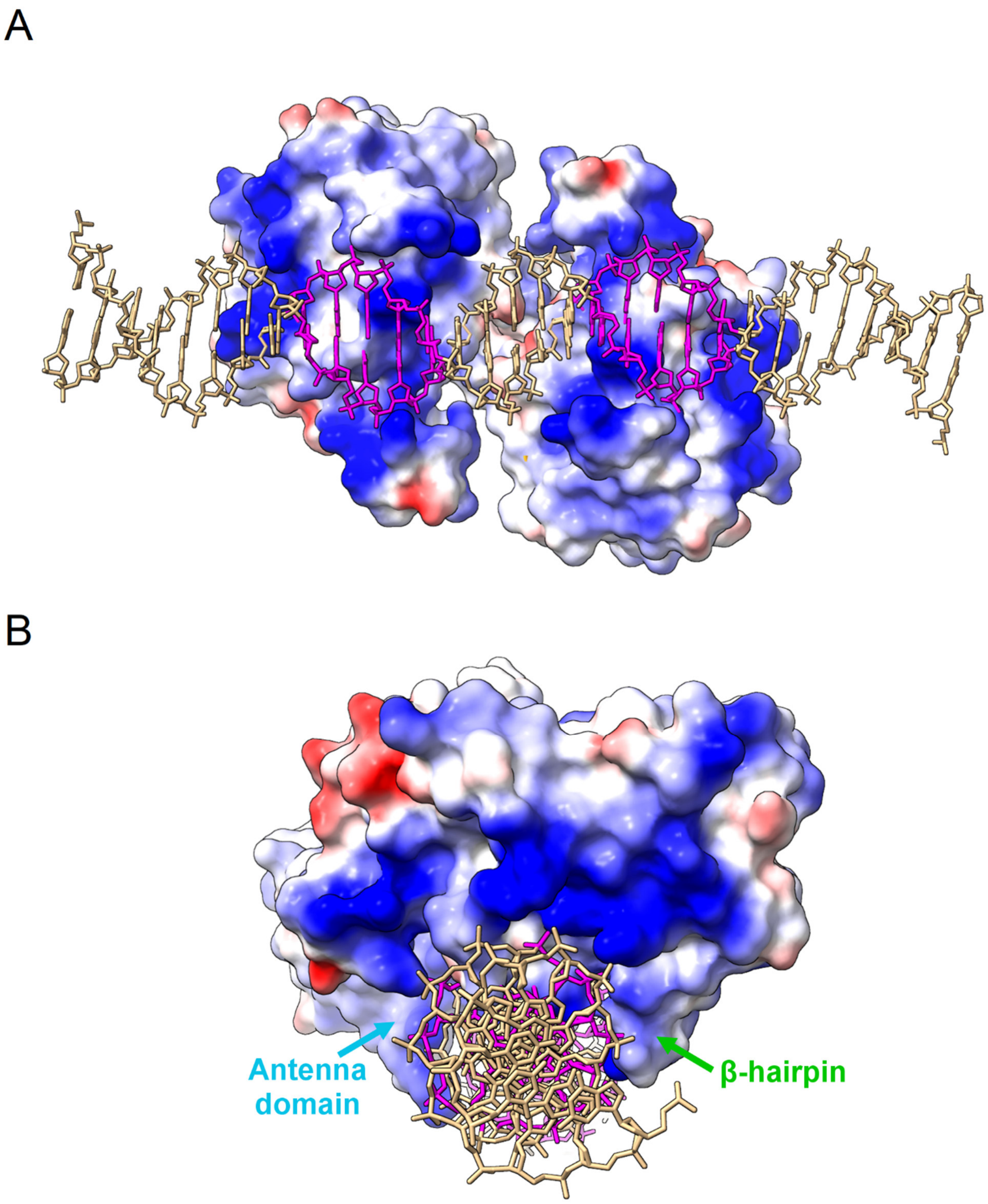


**Fig. S8. Electrostatic surface representation of NFI-X$^{193}$ in complex with DNA. (A)** Bottom view and **(B)** frontal view (similar to Fig. 4C). Red and blue coloring indicate negative and positive regions, respectively. The DNA molecule is shown in stick representation (light brown color), with the palindromic TTGGC(n5)GCCAA NFI-binding site in magenta. This figure was generated using Chimera X version 1.8 [50].

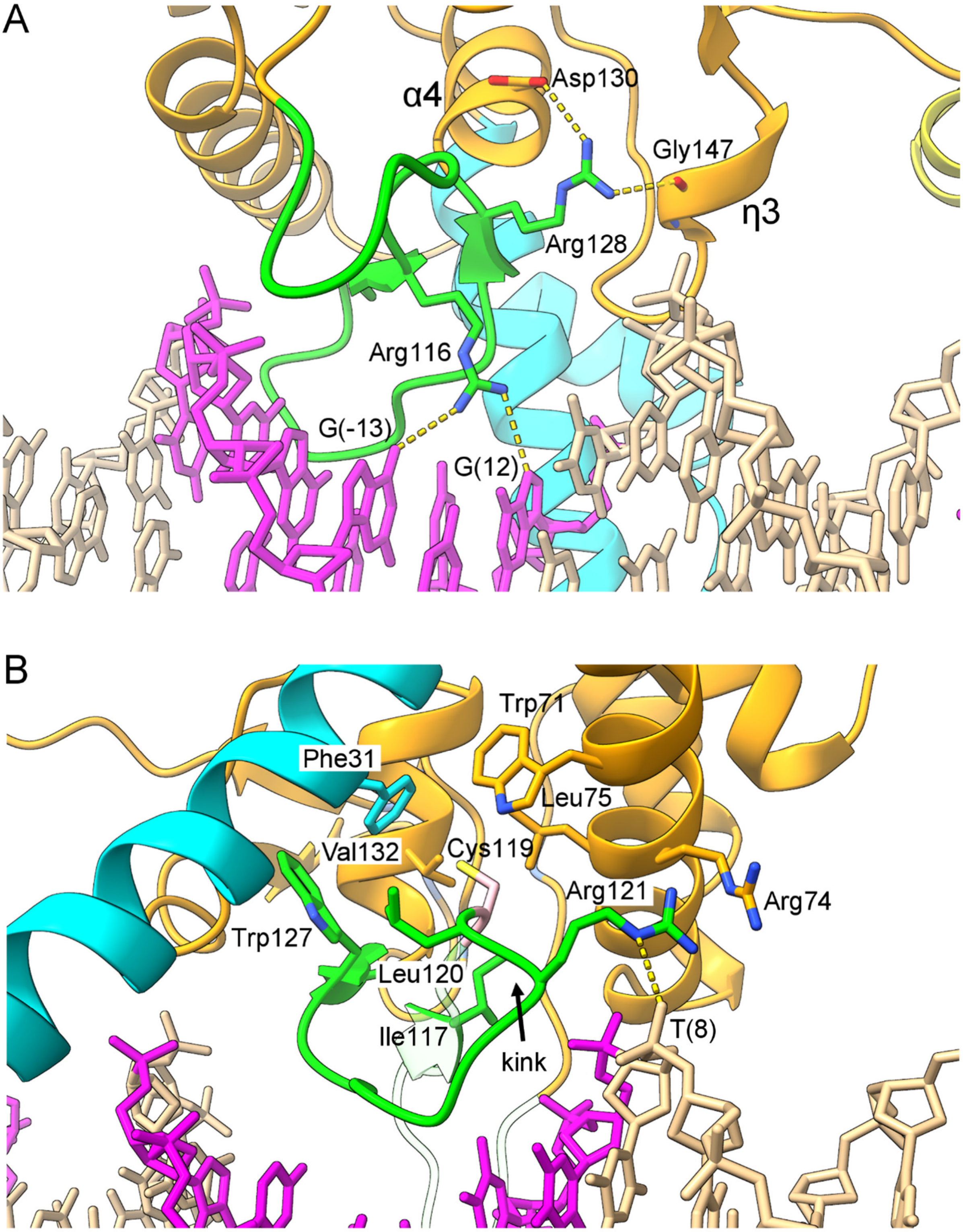


**Fig. S9. -Structural position and interactions of Malan Syndrome residues Arg128 and Arg121.** The structural position and interactions of residues Arg128 and Arg121, found to be mutated in some Malan syndrome individuals (Arg28Leu, Arg121Pro) [6] are shown in panels **(A)** and **(B)**, respectively. Relevant, neighbouring residues are shown in stick representation and labelled, with color coding in accordance with their location in the structure (see Fig. 2A and Fig. S1). The backbone kink at residue Arg128 is indicated. This figure was generated using Chimera X version 1.8 [50].

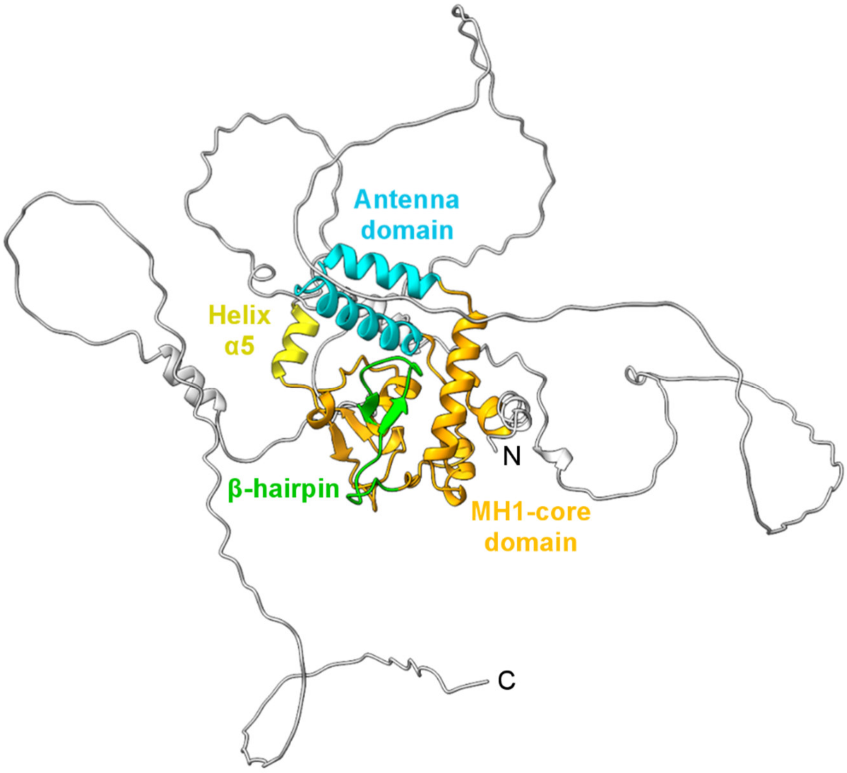


**Fig. S10. AlphaFold prediction of full-length NFI-X.** The NFI-X DBD domains (MH1 and antenna) and regions important for DNA-binding (β-hairpin) and dimerization (α5) are labelled and shown with color coding used in Figure 4A. The C-terminal TAD is shown in grey. The N- and C-terminus are labelled. This figure was generated using Chimera X version 1.8 [50].

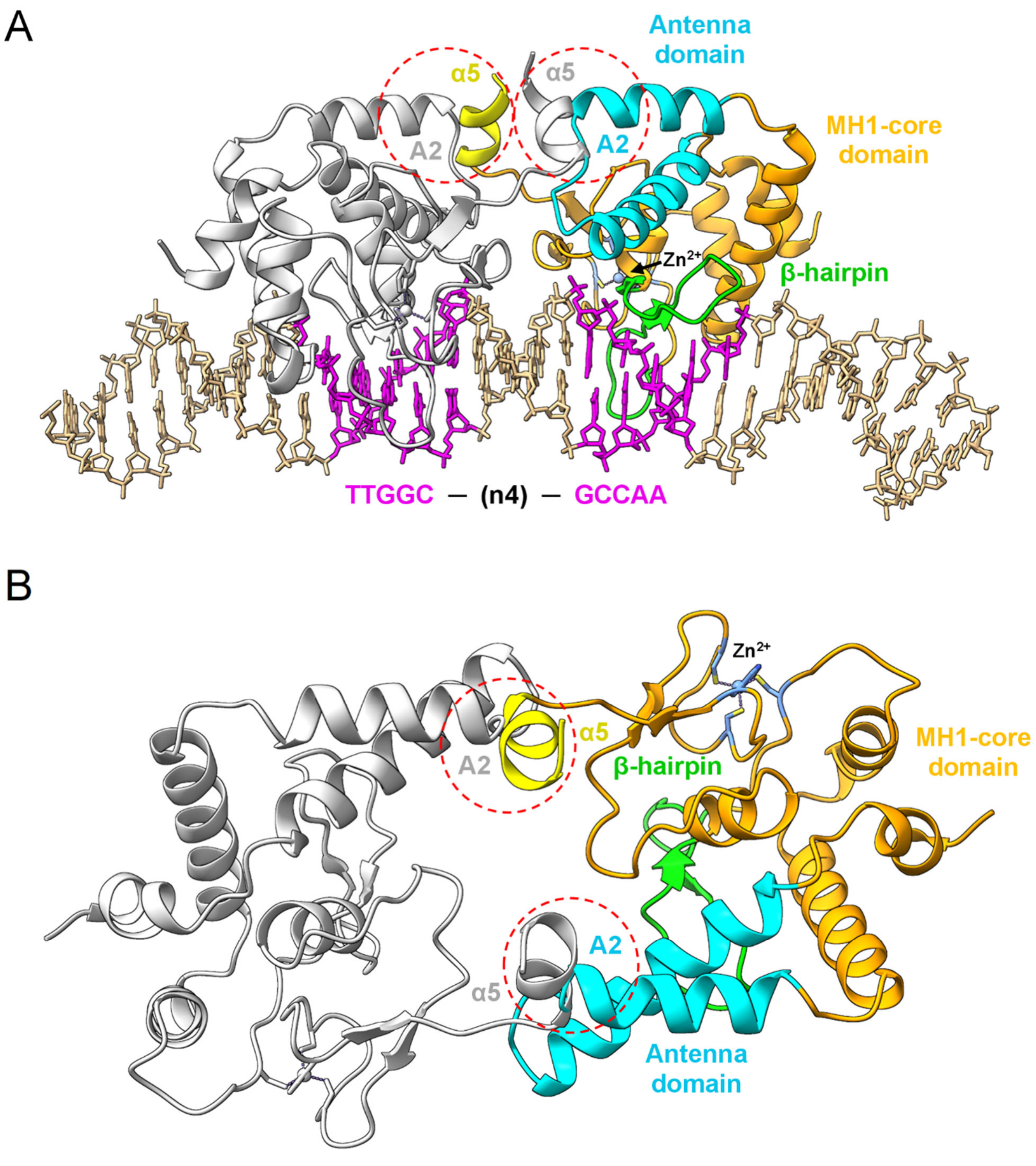


**Fig. S11. Model of the NFI-X$^{193}$ dimer bound to NFI consensus sequence with a shorter, 4bp-spacer. (A)** Side view: one NFI-X$^{193}$ monomer is shown with coloring as in Figure 2, while the other monomer is shown in grey. The main structural regions of the protein are labelled. The DNA molecule is shown in stick representation (light brown coloring), with the palindromic TTGGC(n4)GCCAA NFI-binding site in magenta (as in Fig. 4). Steric clashes between the dimerization helix α5 of one monomer and the A2 helix of the other monomer are highlighted by red dashed circles. (**B**) Top view: for clarity, the DNA molecule has been removed. This figure was generated using Chimera X version 1.8 [50].
